# Lrrns define a visual circuit underlying brightness and contrast perception

**DOI:** 10.1101/2025.05.05.652281

**Authors:** Elena Putti, Giulia Faini, Julie Thanh-Mai Dang, Fanny Eggeler, Jay H. Savaliya, Karine Duroure, Juliette Vougny, Armin Bahl, Jeff W. Lichtman, Florian Engert, Jonathan Boulanger-Weill, Filippo Del Bene, Shahad Albadri

## Abstract

Brightness and contrast are fundamental features of vision, crucial for object detection, environmental navigation, and feeding. Here, we identify a brightness-and contrast-processing circuit in the zebrafish visual system and uncover the role of Leucine-rich repeat neuronal (Lrrn) cell adhesion molecules (CAMs) in regulating its assembly. Deep-projecting retinal ganglion cells (RGCs) serve as the first synaptic relay to the brain requiring Lrrn2 and Lrrn3a for precise axonal targeting and connectivity within the optic tectum. Genetic targeting of these CAMs leads to circuit disorganization and impairments in contrast sensitivity, leading to deficits in visually guided behaviors. Additionally, ultrastructural circuit reconstruction and functional imaging analysis revealed their critical role in luminance processing. These studies define a fundamental visual processing pathway and establish Lrrn CAMs as essential molecular drivers of its assembly.

## Main Text

Alterations in the formation of precise neuronal circuits profoundly impact nervous system function and subsequent behaviors, such as navigation and feeding that rely on brightness and contrast perception and integration (*1–4*). Yet, the developmental and molecular mechanisms governing their assembly are only partially understood. While diverse regulatory mechanisms— including transcription (*5–7*), axon guidance cues (*8–10*), and neuronal activity (*11*) —contribute to circuit formation, synaptic partner recognition is essential for neurons to establish precise connections (*12–16*). This final step in connectivity strongly relies on CAMs (*14*, *17–20*).

We used the larval zebrafish, as an optical and genetically tractable vertebrate model, to characterize a novel neuronal circuit within the visual system (*21*). Our findings reveal that this circuit is essential for processing brightness and/or contrast, key aspects of visual perception (*22*, *23*), and is centered on the *stratum album centrale/stratum periventriculare (*SAC/SPV)-projecting RGCs, deepest layer of the optic tectum (OT), which serves as the first relay of visual information to the brain. RGC axons project from the zebrafish retina to the OT, forming synapses in ten distinct laminae within the superficial neuropil (*24*, *25*). Among these, the SAC/SPV lamina has remained poorly characterized due to imaging and genetic targeting limitations (*26*, *27*). Using a novel RGC transgenic reporter enriched in this lamina *(Tg(gfi1ab:gal4))*, we identified Lrrn CAMs as critical regulators of SAC/SPV-projecting RGC connectivity.

Lrrn proteins belong to the leucine-rich repeat (LRR) family known for their roles, among others, in axon guidance (*8*, *9*), target selection (*5*, *28*, *29*), synapse formation (*30*, *31*), and their involvement in neurological disorders (*32–34*). In *Drosophila*, the Lrrn homolog Capricious (Caps) mediates photoreceptor targeting (*5*). In zebrafish, we show that four Lrrn family members are expressed in the retina, of which Lrrn2, Lrrn3a, and Lrrn3b are selectively expressed in RGCs. Using CRISPR/Cas9-mediated loss-of-function approaches, we demonstrate that Lrrn2 and Lrrn3a CAMs are required for the connectivity of presynaptic SAC/SPV-projecting RGCs with their target neurons both in cell-autonomous and non-autonomous manners. Furthermore, the genetically induced disruption of this circuit leads to severe impairments in brightness and contrast processing, directly linking molecular mechanisms of circuit formation to visual function and behavior. By assigning a novel role to SAC/SPV-projecting RGCs, mapping their synaptic targets through EM segmentation, and integrating genetic, anatomical, functional, and behavioral analyses, our study provides new insights into the genetic control of visual circuit assembly.

### Shedding light onto deep-projecting RGCs

Although deep-targeting RGCs were identified decades ago, they remain poorly characterized, in part because available pan-RGC transgenic lines do not efficiently label them (*35*). In addition, precise characterization of tectal laminae remains challenging due to the dense and overlapping projections of RGCs (*26*, *27*). To target specific subsets of these neurons, we sought to leverage the transcription factor Growth Factor Independent 1ab - *gfi1ab*, which is selectively expressed in a subset of RGCs (*36*) (Supplementary Fig. 1A). Based on this specificity, we generated a novel transgenic reporter line, *Tg(gfi1ab:gal4*) (see Methods). The resulting line, when crossed with *Tg(UAS:lynRFP)*, faithfully reflected the endogenous *gfi1ab* sparse expression in RGCs, as well as in the pineal gland and ear hair cells, consistent with previous findings (*36*) (Supplementary Fig. 1A-B).

We then crossed *Tg(gfi1ab:gal;UAS:lynRFP)* fish with the pan-RGC marker *Tg(isl2b:GFP)* to analyze the projection profiles of *gfi1ab*-expressing RGCs in the retina (Fig. 1A) and OT (Fig. 1A^I^) of developing larvae. Although *gfi1ab* did not label a specific morpho-group of RGCs, it is expressed in a salt-and-pepper manner across different subpopulations (Supplementary Fig. 1C-D). These Gfi1ab-RGC axons projected to all OT layers, including SO, SFGS, SGC, and particularly in the SAC/SPV layer (Fig. 1 A^I^). This unique enrichment of *gfi1ab* expression in SAC/SPV-projecting RGCs provides a novel tool for labeling deep-projecting axons in this tectal lamina that complements pan-RGC transgenic lines such as Tg(*isl2b:GFP*), where the strong fluorescence from superficial layers (SO/SFGS) makes challenging functionally imaging deeper projections (Supplementary Fig. 1E-F).

**Fig. 1.**
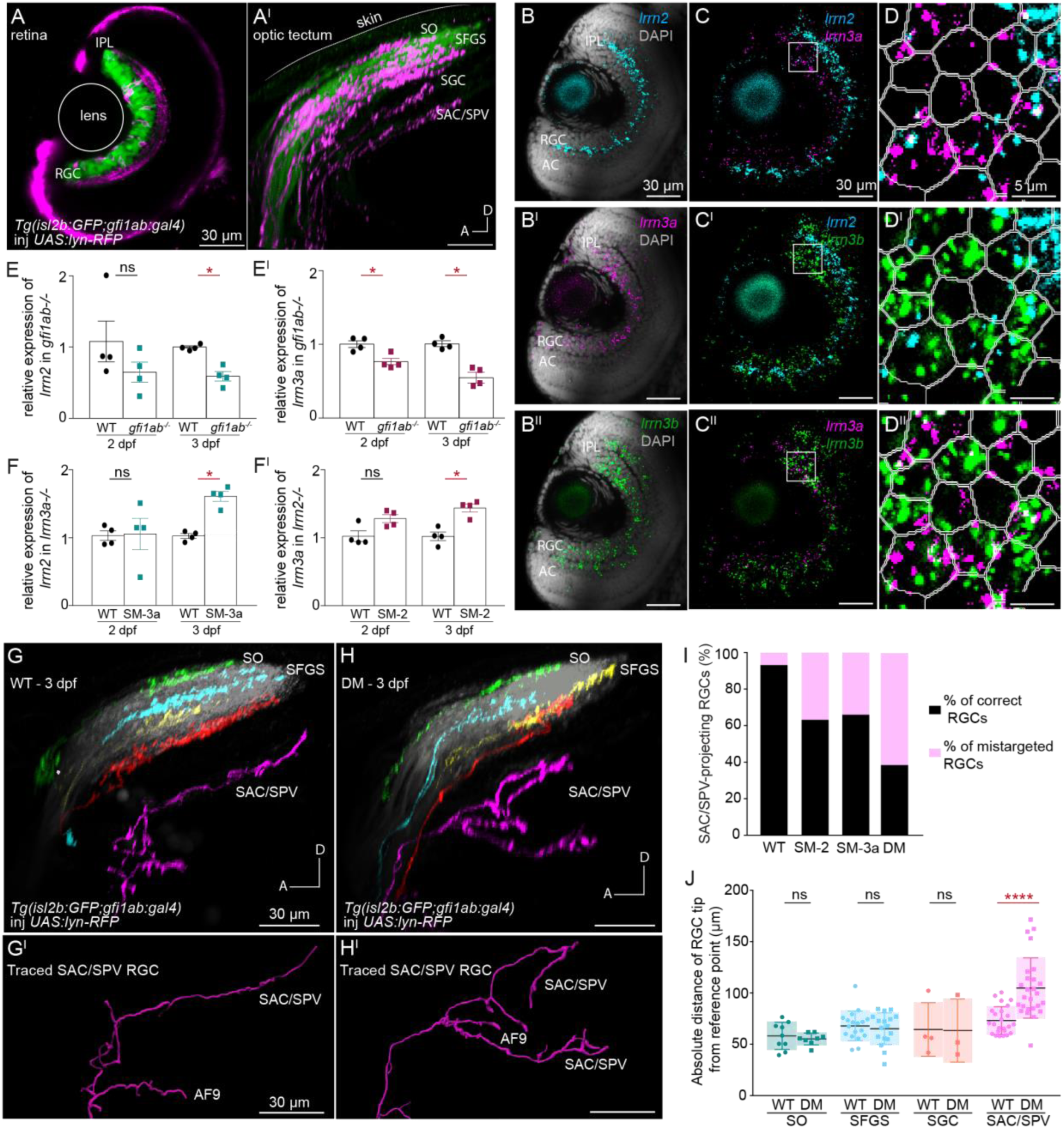
Gfi1ab and Lrrns are expressed in the developing RGCs of the zebrafish retina and are required for RGC wiring assembly. (A-A^I^) Confocal images of 3 dpf *Tg(gfi1ab:gal4;isl2b:GFP)* embryos injected at the 1-cell stage with a *10UAS:lynRFP* plasmid. (A) *Gfi1ab:gal4* expression (magenta) is sparse in RGCs compared to the pan-RGC marker Isl2b (green). (A^I^) *Gfi1ab*-RGC axons project within all sub-laminae of the tectal neuropil. (B-D^II^) Confocal Z-stack images of a 3 dpf retina stained by multiplex HCR *in situ* hybridization for *lrrn2* (cyan), *lrrn3a* (magenta), and *lrrn3b* (green) showing their relative expression in the GCL and ACs. DAPI (gray) highlights retinal layers. (B-B^II^) Single-channel expression of *lrrn2* (cyan), *lrrn3a* (magenta), and *lrrn3b* (green). (C-C^II^) Partial colocalization of *lrrn2/lrrn3a*, *lrrn2/lrrn3b*, and *lrrn3a/lrrn3b*. (D-D^II^) Magnified images using Cellpose to identify colocalization within RGCs. (E-E) Quantitative RT-PCR analysets of *lrrn* isoforms expression in *gfi1ab-/-* mutants at 2 and 3 dpf. Significant downregulation of *lrrn2* (*p* = 0.0286) (E) and *lrrn3a* (E’) is observed at 3 dpf (*p* = 0.0286), with only *lrrn3a* down-regulated at 2 dpf (*p* = 0.0286). Data are mean ± SD; n = 3 technical replicates, 50 heads per condition, n = 4 biological replicates. (F-F^I^) Quantitative RT-PCR analyses of *lrrn* isoforms expression in single mutant *lrrn2* (SM-2) and single mutant *lrrn3a* (SM-3a) mutants at 2 and 3 dpf. *Lrrn3a* is upregulated in SM-2 at 3 dpf (*p* = 0.0286) (F), and *lrrn2* is upregulated in SM-3a at both 2 and 3 dpf (*p* = 0.0286) F’. Data are mean ± SD; n = 3 technical replicates, 50 heads per condition, n = 4 biological replicates. (G-H^I^) Confocal imaging of axonal projections of 3 dpf WT (G) and *lrrn2^-/-^; lrrn3a^-/-^*double mutant (DM) embryos (H) Gfi1ab-RGCs in the *Tg(gfi1ab:gal4;isl2b:GFP)* background injected with a *10UAS:lynRFP* construct for mosaic labeling of Gfi1ab-RGC axonal projections. WT Gfi1ab-RFP positive RGCs target unique tectal laminae, whereas DM Gfi1ab-RFP positive RGCs show mistargeting in the deep SAC/SPV region. (G^II^, H^II^) Magnified views of SAC/SPV-RGC axons. (I) Percentage of correctly targeted and mistargeted Gfi1ab-RGCs in WT, DM, and SM embryos. (J) Quantification of the average absolute distances (μm) of RGC axonal tips from the skin for panels (G) and (H), showing a significant difference in the SAC/SPV layer of DM larvae compared to WT. Data are mean ± SD. *p* < 0.0001. Scale bars = 30 μm, unless stated otherwise. Statistical significance was determined by a Mann-Whitney U-test: ns – *p* > 0.05; \**p* < 0.05; \*\**p* < 0.01; \*\*\**p* < 0.001; \*\*\*\**p* < 0.0001. Abbreviations: AC = amacrine cells; GCL = ganglion cell layer; IPL = inner plexiform layer; OT = optic tectum; RGC = retinal ganglion cells; SAC = stratum album centrale; SFGS = stratum fibrosum et griseum superficiale; SGC = stratum griseum centrale; SO = stratum opticum; SPV = stratum periventriculare; AF9 = arborization field 9; WT = wild-type; DM = double mutant (*lrrn2^-/-^; lrrn3a^-/-^);* SM-2 = single mutant *lrrn2^-/-^*; SM-3a = single mutant *lrrn3a^-/-^*.

In *Drosophila*, the ortholog of zebrafish Gfi1ab transcription factor, Senseless, regulates the expression of the LRR CAM Caps, which plays a key role in photoreceptor targeting (*5*, *28*). We assessed the conservation of this pathway in vertebrates by performing an *in silico* search analysis (Homologene NCBI) of Caps orthologues. We identified the Lrrn CAM protein family, comprising *lrrn1, lrrn2, lrrn3a,* and *lrrn3b*. Whole-mount *in situ* hybridization at 2 and 3 days post fertilization (dpf) revealed early and broad expression of all four paralogues in the fish’s forebrain, midbrain, and hindbrain (Supplementary Fig. 2A-B). Retinal cross-sections at 3 dpf showed *lrrn2, lrrn3a,* and *lrrn3b* expression in RGCs, with *lrrn2* and *lrrn3b* also present in the amacrine cell (AC) layer (Supplementary Fig. 1 G-G^I^). Multiplex HCR combined with confocal imaging showed that the expression of these three *lrrn* members partially overlap in the GCL (Fig. 1B-D). Using Cellpose (*37*), we extracted the nuclear DAPI-derived signal delineated the RGC membrane borders and observed that the HCR signals colocalized within the same cells (Fig. 1D).

### Gfi1ab and Lrrn genetic deletions lead to mistargeting defects in the deep-OT

To investigate the roles of Gfi1ab and Lrrn CAMs in vertebrates for RGC projections, we adopted a loss-of-function approach in zebrafish using CRISPR/Cas9 mediated KOs to either introduce early stop codons (*gfi1ab^-/-^)* or disrupt the signal peptide and the LRR domains (*lrrn^-/-^*) (see Methods) (Supplementary Fig. 2A). All mutant lines generated were viable at homozygosity and showed no gross morphological defects.

We first assessed the impact of *gfi1ab* loss of function on *lrrn* CAMs expression levels in the larval visual system. To do so, we performed qRT-PCR on the heads of *gfi1ab*^-/-^ mutants and wild type (WT) controls at 2 and 3 dpf. *Lrrn2* and *lrrn3a* mRNAs were significantly downregulated in the absence of Gf1ab, particularly at 3 dpf (Fig. 1E, Supplementary Fig. 2B), while *lrrn3b* and *lrrn1* levels remained unchanged (Supplementary Fig. 2C-C^I^). HCR experiments performed in 3 dpf *Tg(gfi1ab:gal4;UAS:lyn-GFP)* larvae showed significant co-localization of *lrrn2* (Supplementary Fig. 2D) and *lrrn3a* (Supplementary Fig. 2DI) within *gfi1ab-*expressing cells. These observations further pinpoint a shared molecular pathway. In addition, qRT-PCR on the heads of *lrrn2* and *lrrn3a* mutant larvae performed to assess expression changes in the other CAMs part of the *lrrn* subfamily, indeed revealed a compensatory mechanism between both *lrrn2* and *lrrn3a* specifically. *Lrrn2* was upregulated in *lrrn3a*^-/-^ mutants (SM-3a) at 3 dpf (Fig. 1F) and similarly, *lrrn3a* was upregulated in *lrrn2^-/-^* mutants (SM-2) (Fig. 1F^I^). Instead, *lrrn3b* and *lrrn1* remain unchanged in both mutants. (Supplementary Fig. 2E-E^I^).

Next, to evaluate the impact of the loss of Gfi1ab on *gfi1ab*-expressing RGC development, we imaged *Tg*(*gfi1ab:gal4;UAS:mRFP*)*;gfi1ab*^-/-^ larvae at 3 dpf when all RGCs have projected their axons to the OT. However, unlike WT control larvae, no fluorescence could be detected in the *gfi1ab* mutant (Supplementary Fig. 2F). A Tunel assay on *gfi1ab*^-/-^ larvae revealed no increase in apoptosis compared to controls (Supplementary Fig. 2G), suggesting a self-regulatory mechanism of Gfi1ab on its promoter that is disrupted in Gfi1ab mutants. With *lrrn2* and *lrrn3a* downregulated in *gfi1ab*^-/-^ mutants, we evaluated the impact of their loss of functions as well as their combined depletion on *gfi1ab*-expressing RGC projections in the OT. We generated single (SM) and double mutant (DM) lines in the *Tg(gfi1ab:gal4;isl2b:GFP)* transgenic background and injected a *10UAS:lynRFP* plasmid into SM-2, SM-3a, and DM larvae at the 1-cell stage, enabling single-cell analysis by mosaically labeling RGC membranes and axons. The *Tg(isl2b:GFP)* background served as a reference to identify precise tectal laminae. While *lrrn* mutants did not show any obvious retinal or optic tract defects (Supplementary Fig. 2H-I), SM-2, SM-3a, and DM larvae exhibited clear axonal mistargeting precisely in the SAC/SPV layer (Fig. 1G-H). Compared to WT larvae, where RFP-positive RGCs projected to single and precise laminae of the OT, over 25% of SAC/SPV-projecting RGCs deviated from their normal target in the SM-2 and SM-3a larvae (Fig. 1I). DM larvae displayed even more severe defects than SM larvae, with >60% of deep-projecting RGCs exhibiting miswiring (Fig. 1I), suggesting an additive effect of *lrrn2* and *lrrn3a* losses. Tracing the different RGCs within the OT revealed no significant differences in RGCs projecting to the SO, SFGS, and SGC layers between DM larvae and WT controls (Fig. 1J). However, RGCs projecting to the SAC/SPV layer in DM larvae showed a significantly increased distance from the reference point on the skin compared to WT controls (Fig. 1J). These defects persisted at 5 dpf (Supplementary Fig. 3E), indicating that Lrrn2 and Lrrn3a are key CAM regulators of SAC/SPV targeting. Similar SAC/SPV-related miswiring defects were also detected in the *gfi1ab^-/-^* larvae at 3 dpf injected with the *isl2b:gfp* plasmid at the 1-cell stage (Supplementary Fig. 3F).

Finally, we also investigated whether *lrrn3b,* also expressed in RGC, would induce defects in the axonal projections of Gfi1ab-RGCs when knocked out (KO). We thereby injected a *UAS:GFP* construct at 1-cell stage to label *gfi1ab*-expressing RGCs in SM-2, SM-3a and SM-3b larvae.

Compared to SM-2 and SM-3a, no defects were observed in SM-3b larvae whose phenotype appeared to be reminiscent of those of WT control larvae (Supplementary Fig. 3A-D). Taken together, our results demonstrate that Lrrn2 and Lrrn3a function downstream of Gfi1ab in the vertebrate visual system, contributing to the accurate targeting of *gfi1ab*-expressing deep-targeting RGCs to the SAC/SPV layer of the OT.

### Deep-targeting SAC/SPV RGCs molecular requirements for proper axonal projection

We then asked whether, at the cellular level, these CAMs are required within RGCs to achieve proper tectal lamination or act via non-cell-autonomous mechanisms. To address this, we performed cell transplantation assays at the blastula stage. First, we transplanted DM RGCs lacking Lrrn2 and Lrrn3a into a WT environment to determine if defects persisted in a WT OT (Fig. 2A). Chimeric embryos were generated by transplanting donor cells from *10UAS:lynRFP*-injected WT or DM *Tg(gfi1ab:gal4)* embryos into WT *Tg(isl2b:GFP)* hosts of the same developmental stage (Fig. 2A). Confocal imaging of the resulting chimeric larvae at 3 dpf was performed to assess the axonal development of WT or DM *gfi1ab*-expressing RGCs devoid of Lrrn2 and Lrrn3a transplanted into WT. Our analysis revealed that mistargeting defects persisted when DM cells developed in a WT environment, revealing a cell-autonomous requirement of Lrrn2 and Lrrn3a in RGC targeting (46%) (Fig. 2D-E). Next, we assessed the contribution of the OT by transplanting WT RGCs (expressing Lrrn2 and Lrrn3a) into a DM host (Fig. 2F). Over 44% of deep-projecting RGCs expressing both CAMs failed to reach their targets in the DM OT, differently from controls (Fig. 2G-J). These results indicate that Lrrns are essential for proper axonal targeting both in the pre-synaptic retina and the post-synaptic partners in the OT. Having established that Lrrns function in both pre-and post-synaptic compartments to regulate RGC targeting, we next sought to unravel the molecular mechanisms underlying their role in axon guidance. In *Drosophila*, Caps is thought to mediate targeting through homophilic binding interactions (*5*). To determine whether zebrafish Lrrns operate through similar or distinct mechanisms, we performed an *in vitro* protein aggregation assay to assess their binding properties. We asked whether, like Caps, Lrrn CAMs act through homophilic or heterophilic interaction or both. We performed an *in vitro* protein aggregation assay using HEK293 cells transfected with *pCX:lrrn2-GFP* or *pCX:lrrn3a-RFP*, with *pCX:GFP* and *pCX:RFP* as controls (Fig. 2K-O, Supplementary Fig. 4A-D). To evaluate homophilic interaction, cells expressing either Lrrn2-GFP or Lrrn3a-RFP showed an increase in aggregation compared to controls (Fig. 2K-L, 2N, Supplementary Fig. E-F^I^). To assess if both CAMs could interact through heterophilic binding, Lrrn2-GFP and Lrrn3a-RFP cells were co-aggregated. We observed a higher number of aggregates co-expressing both fluorophores (Fig. 2M, O, Supplementary Fig. G-H). Together these results indicate that Lrrn2 and Lrrn3a can bind through both homophilic and heterophilic interactions.

**Fig. 2.**
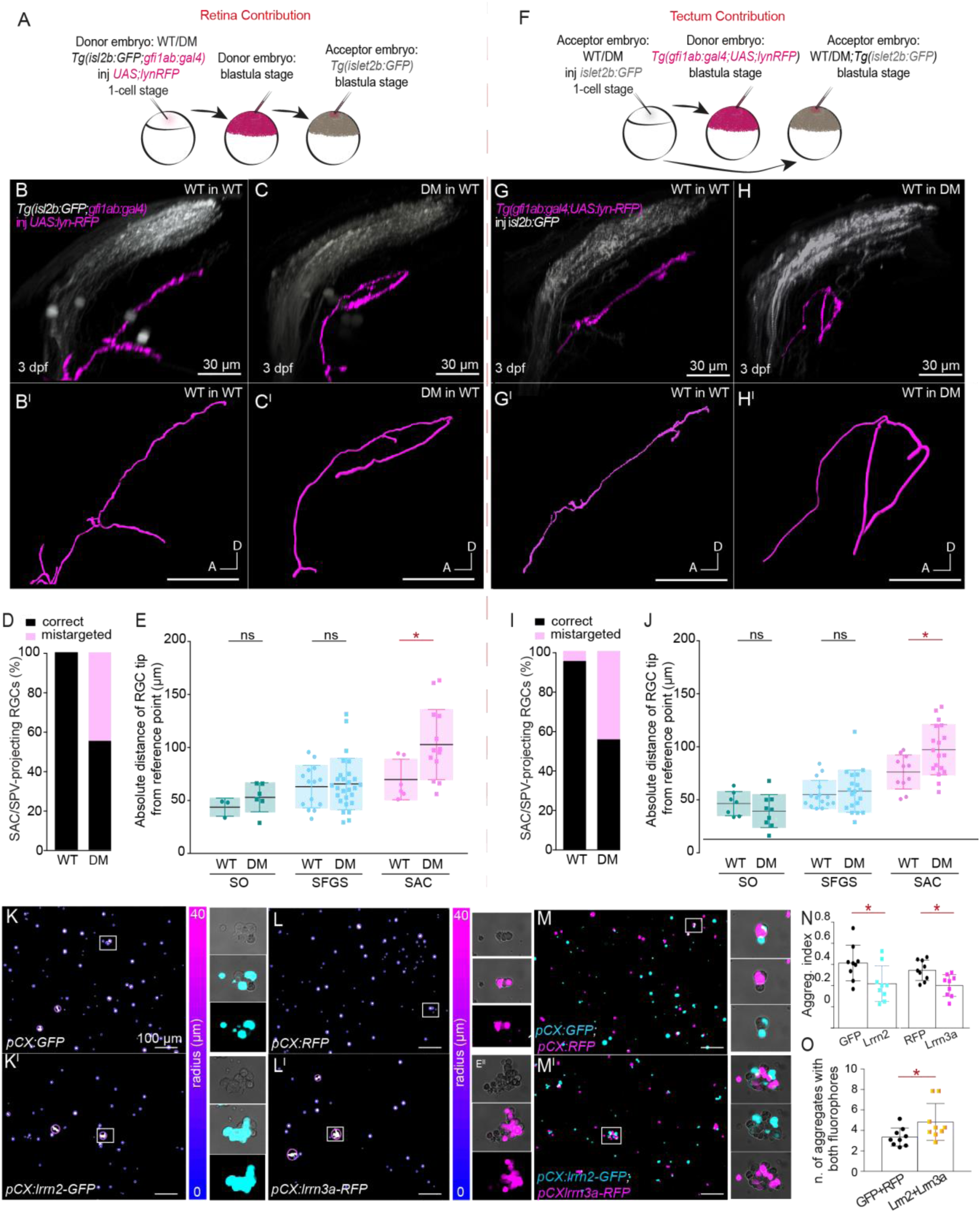
Lrrn2/Lrrn3a are required both in the retina and in the optic tectum for the proper projection targeting of Gfi1ab-RGC and act through homophilic and heterophilic interactions. (A, **F)** Transplantation experimental workflows at blastula stage for assessing retina (A) and tectal (F) contributions of Lrrn2 and Lrrn3a for Gf1ab RGCs axonal projections in 3 dpf chimeric zebrafish embryos. **(B-C)** Confocal images at 3 dpf of chimeras with WT (**B**) and DM (C) *gfi1ab*-RGCs transplanted into WT embryos, showing SAC/SPV projections and mis-projections (B-I). (B^I^) and (C^I^) panels show magnified views of Gfi1ab-RFP positive SAC/SPV-RGC axon projections. **(D)** Percentage of correctly targeted and mistargeted SAC/SPV-RGCs in WT and DM transplanted into WT embryos (WT = 6, DM = 13 SAC/SPV-RGCs). **(E)** Quantification of axonal distances from the skin for panels (B) and (C) showing significant difference for the SAC/SPV layer, increased for DM compared to WT SAC/SPV-RGCs *p* = 0.0365. **(G-H)** Confocal images showing WT *gfi1ab*-RGCs transplanted in WT (G) and DM embryos (H), highlighting SAC/SPV projections at 3 dpf. (G^I^) and (H^I^) panels show magnified views of SAC/SPV-RGC axon projections. **(I)** Percentage of correctly targeted and mistargeted SAC/SPV-RGCs in WT and DM transplanted into WT embryos (WT = 12, DM = 18 SAC/SPV-RGCs). **(J)** Quantification of axonal distances for WT and DM transplanted into WT embryos with significant difference for the SAC/SPV layer increased, for DM compared to WT SAC/SPV-RGCs (*p* = 0.0311). **(K-L)** Aggregation assay of HEK293 cells transfected with *pCX:GFP*. (K) or *pCX:RFP* (L) control plasmids. (K^I^-L^I^) Aggregation assay of HEK293 cells transfected with *pCX:lrrn2-GFP* (K^I^) or *pCX:lrrn3a-RFP* (L^I^), showing the largest aggregates. **(M-M^I^)** Co-aggregation assays of HEK293 cells transfected with *pCX:GFP* and *pCX:RFP* (M), or with *pCX:lrrn2-GFP* and *pCX:lrrn3a-RFP* (M^I^), to assess heterophilic interactions. **(N)** Aggregation index for control and Lrrn2/Lrrn3a overexpression conditions. Lrrn2: *p* = 0.04; Lrrn3a: *p* = 0.0306. **(J)** Number of aggregates co-expressing both GFP (cyan) /RFP (magenta) and Lrrn2 (cyan) /Lrrn3a (magenta). Statistical analyses: Non-parametric Mann-Whitney U-test. Data represented as mean ± SD. \**p* < 0.05. (B-H) Scale bar = 30 μm. (K-M) Scale bar = 100 μm. Dotted lines separate retina and tectum contributions. Abbreviations:SFGS, stratum fibrosum et griseum superficiale; SGC, stratum griseum centrale; SO, stratum opticum; SPV, stratum periventriculare; WT, wild-type.

### SAC/SPV-projecting RGCs in the visual system are sensitive to luminance changes

Previous studies have shown that both neurons from the periventricular zone (PVZ) and non-tectal neurons can project to the deepest layers of the OT, but a comprehensive connectome remains unresolved (*26*, *38–42*). To investigate this, we performed an ultrastructural analysis using a novel electron microscopy (EM) dataset, combining two-photon imaging of 7 dpf zebrafish with EM volume data to generate a functional connectome, achieving single-cell precision in visualizing neural structures (*43*). This dataset enabled us to map the detailed morphology and synaptic connections of SAC/SPV axons in the lower half of the OT (Supplementary Fig. 5A). We identified SAC/SPV RGC axons, and due to its high-quality resolution, we could reconstruct and distinguish synaptic density regions, allowing us to detect axo-dendritic, axo-axonic, and axo-somatic synapses (Fig. 3A-B, Supplementary Fig. 5B). For all the four traced SAC/SPV-RGCs, most synapses were axo-dendritic. Notably, we also identified previously undetected synapse types: axo-axonic and axo-somatic, which were not visible using classical light microscopy (Supplementary Fig. 5B). Further analysis of the axo-dendritic connections revealed that they primarily targeted non-stratified periventricular interneurons (nsPVINs) and projection neurons (nsPVPNs), making them the main postsynaptic targets of SAC/SPV projecting RGCs (Fig. 3C-D, Supplementary Fig. 5C-D). Some connections also involved other PVN classes, including inhibitory PVNs (Fig. 3D). Nonetheless, we found that the majority of SAC/SPV postsynaptic targets are excitatory neurons (Supplementary Fig. 5E). Given that *lrrn2* and *lrrn3a* are expressed in the PVZ, and that these nsPVINs and nsPVPNs are key targets of SAC/SPV axons, we wondered whether these PVNs might be affected in our mutants. To address this question, we injected the *foxp2:gal4* plasmid and analyzed PVNs morphology in DM larvae, as previously performed (*44*). However, we did not find any difference in the proportion of stratified (sPVNs, which include mono, bi and tri stratified PVPN) versus non-stratified PVNs (nsPVNs, which include both nsPVPNs and nsPVINs) between WT and DM larvae, nor did we observe any clear mistargeting defect (Fig. 3E, Supplementary Fig. 5F), suggesting that Lrrn2 and Lrrn3a are not directly required for the development of these cells.

**Fig. 3.**
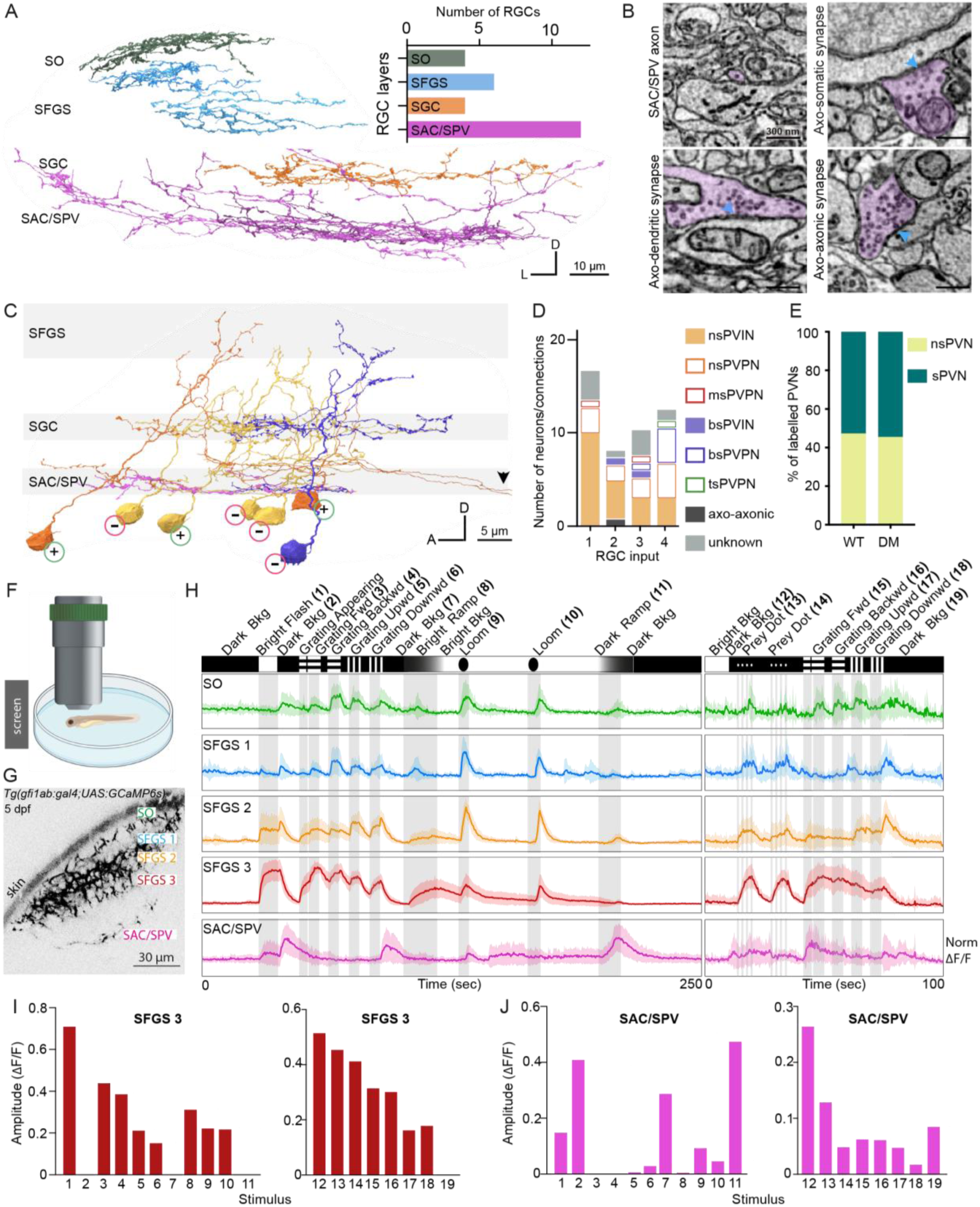
Synaptic Connectivity and Functional Properties of SAC/SPV RGCs in the Optic Tectum. **(A)** Dorsal view of reconstructed RGCs spanning all layers of the left optic tectum (OT) from the serial electron microscopy (SEM) volume dataset. The histogram shows the number of reconstructed RGC per layer. **(B)** Types of synapses identified within a SAC/SPV axon: axo-dendritic, axo-somatic, and axo-axonic. (**C)** Sparse connectome of an example SAC/SPV axon (pink), displaying the most represented post-synaptic targets identified within the OT volume. The most superficial OT layers are not shown as none of the post-synaptic neurons innervated them. Arrows show axons of PVPN neurons leaving the OT. **(D)** Bar plot showing the distribution of identified PVN types within 4 SAC/SPV-RGCs, with color-coded PVN classes matching between panels. Filled boxes correspond to interneurons while hollow boxes correspond to projection neurons. RGC n. 2 corresponds to sparse connectome shown in (C). Color-coding is maintained between (C) and (D). **(E)** Bar plot showing the distribution in percentage (%) of sPVNs versus nsPVNs in DM and WT controls. (n = 38 PVNs, from WT 27 larvae, n = 33 PVNs from DM = 21 larvae). **(F)** Illustration of the setup used for functional calcium imaging in *in vivo* 5-6 dpf larvae, with visual stimuli projected in front of the fish’s eye. **(G)** Two-photon image of tectal lamination of a *Tg(gfi1ab:gal4;UAS:GcaMP6s)* 5 dpf embryo, with different regions within the volume discriminated. Scale bar = 30 μm. **(H)** Functional responses from distinct tectal regions shown in (G) to a series of visual stimuli (illustrated above the traces). Data are averaged across larvae (n = 11 for the first stimulus set; n = 7 for the second). Shaded areas represent standard deviation (SD). **(I)** Quantification of normalized response amplitudes (ΔF/F) for each stimulus in the SFGS 3 region, corresponding to the stimuli shown in (H). **(J)** Quantification of normalized response amplitudes (ΔF/F) for each stimulus in the SAC/SPV region, corresponding to the stimuli shown in (H). Abbreviations: OT, optic tectum, SFGS, stratum fibrosum et griseum superficiale; SGC, stratum griseum centrale; SO, stratum opticum; SPV, stratum periventriculare; PVIN, periventricular interneuron; PVPN, periventricular projection neuron, nsPVIN: non-stratified PVIN; nsPVPN, non-stratified PVPN; ms, mono stratified; bs, bistratified; ts, tristratified; s, stratified; Bkg, background; WT, wild-type; DM, double mutant *lrrn2^-/-^*; *lrrn3a^-/^*^-^.

Previous calcium imaging studies of the OT and pretectum have identified distinct functional groups within RGCs but had limited insights on the functional properties of the SAC/SPV layer (*26*, *27*, *45*). To address this knowledge gap, we used our *Tg(gfi1ab:gal4;UAS:GcaMP6s)* transgenic line. First, we employed a battery of visual stimuli, based on those used in previous studies (*26*, *27*), and successfully subdivided the neuropil into five regions—SO, SFGS1, SFGS2, SFGS3 and SAC/SPV—according to their distance from the skin (Fig. 3F-G).

Functional responses in SO and SFGS regions were consistent with prior findings (*27*), confirming the reliability of the stimulus set (Fig. 3H-I, Supplementary Fig. H-J).

In contrast, in the SAC/SPV layer, we observed a robust activation to luminance changes, particularly when brightness decreased. This was most pronounced during dark and bright flashes (stimuli n.1 and 2 in Fig. 3H), as well as the dark ramp stimulus (n. 11), and was also observed when the stimuli were turned off (e.g. n. 7 and 19) (Fig. 3H, 3J, Supplementary Fig. 5G). To explore more specific responses, we presented prey-like stimuli and gratings independently of the standard stimulus battery. While the SAC/SPV layer showed some activation to prey-like dots (n. 13 and 14), it did not generate a distinct fluorescence peak as seen in the more superficial layers. Notably, its response emerged slightly earlier and persisted longer. A similar prolonged response pattern was observed when gratings were reintroduced (n. 15-18) - an effect that had not been detected during the initial stimulus presentation, possibly because the preceding stimulus (appearance of a dark background) in the original battery elicited strong neuronal activation that masked subsequent responses (Fig. 3H, 3J). Activation persisted even after the gratings were turned off, extending until the background transitioned to black. Together, these findings demonstrate that the SAC/SPV layer is highly sensitive to luminance changes, particularly to brightness decreases, but does not encode direction or orientation.

### Altered SAC/SPV projections lead to impairments in brightness and contrast detection

The pronounced laminar defects observed in DM larvae raised the question of whether these structural abnormalities also led to visual impairments, especially given their distinct functional responses to dimming stimuli (Fig. 3H). To assess this, we first ruled out general motor impairments by tracking freely swimming 6 dpf WT and DM larvae for several minutes, finding no significant differences (Supplementary Fig. 6A-A^III^).

To evaluate visual deficits, we examined hunting behavior and optomotor responses (OMR) in 6 dpf larvae. Initially, a prey-consumption assay was conducted to compare the overall hunting efficiency between DM and WT larvae. The assay revealed a significant reduction in the number of rotifers consumed by the DM larvae during the 4 hours of monitoring compared to WT controls (Fig. 4B), indicating a defect in prey-capture. Similar deficits were observed for SM-2, SM-3a, and *gfi1ab^-/-^* larvae, revealing the involvement of this molecular pathway in the correct development of the neuronal circuits associated with this behavior (Fig. 4C, Supplementary Fig. 4B-B^I^).

**Fig. 4:**
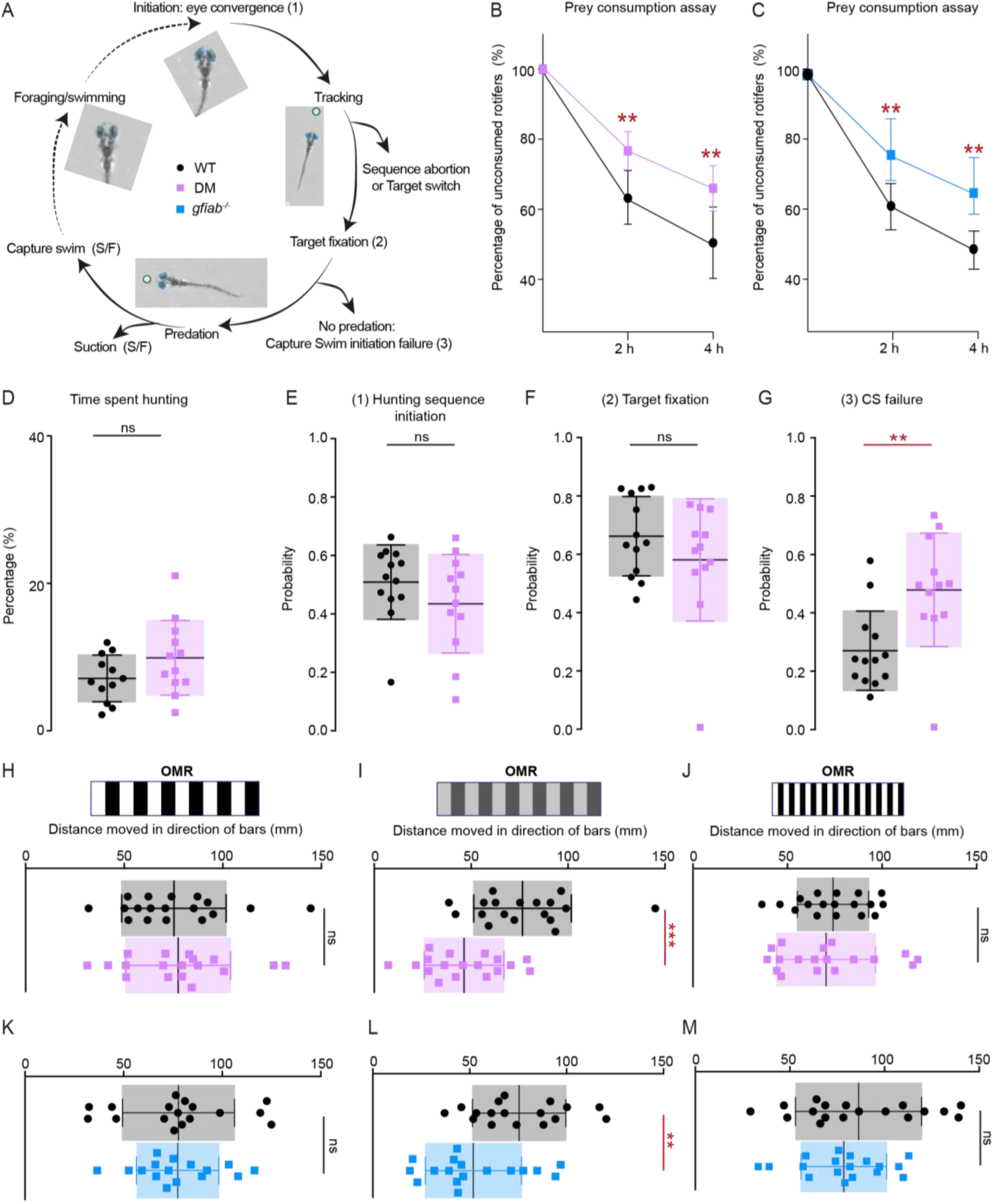
**Lrrn2, Lrrn3a and Gfi1ab loss of functions lead to impairments in visually guided behaviors**. **(A**) Schematic representation of the hunting sequence in 6 dpf WT and DM larvae. Full circle indicates a successful hunting sequence, while outward arrows indicate premature termination or failure. Eyes circled in light blue highlight convergence. (**B-C**) Prey-consumption assay at 6 dpf, showing the percentage of rotifers consumed by WT and DM larvae (B), and WT and *gfi1ab^-/-^* larvae (C) after 2 and 4 hours. DM and *gfi1ab^-/-^* larvae consume significantly fewer rotifers than WT controls (n = 10 larvae per genotype. DM: *p* = 0.0013; *gfi1ab^-/-^ p* = 0.0014). **(D)** Percentage of time spent hunting with converged eyes in 6 dpf WT and DM larvae. No significant difference in hunting time between the groups (number of larvae: WT = 12, DM = 12; *p* > 0.05). **(E-G)** Analysis of hunting sequence events in 6 dpf WT and DM larvae (matching labels in panel A) (number of fish: WT = 12, DM = 12; total hunting sequences: WT = 1084, DM = 1249). **(E)** Probability of initiating hunting sequences: no significant difference between WT and DM larvae. *p* > 0.05. **(F**) Target fixation probability: no significant difference between WT and DM larvae. *p* > 0.05. (**G)** Probability of initiating capture swims: significant impairment in DM larvae compared to WT. *p* = 0.0037. **(H-M)** Optomotor response (OMR) assay with moving bars (left or right). **(H**) Distance swam by WT and DM larvae with 10 mm gratings at 100% contrast difference (number of larvae: WT = 18, DM = 18). **(I)** Distance swam by WT and DM larvae with 10 mm gratings at 20% contrast difference. DM larvae swim significantly less than WT. Number of larvae: WT = 18, DM = 18. *p* = 0.0006. **(J)** Distance swam by WT and DM larvae with 4 mm gratings at 100% contrast difference (number of larvae: WT = 18, DM = 18). **(K)** Distance swam by WT and *gfi1ab^-/-^* larvae with 10 mm gratings at 100% contrast difference (number of larvae: WT = 18, *gfi1ab^-/-^* = 18). **(L)** Distance swam by WT and *gfi1ab^-/-^* larvae with 10 mm gratings at 20% contrast difference. *Gfi1ab^-/-^* larvae show reduced swimming distance compared to WT (number of larvae: WT = 18, *gfi1ab^-/-^* = 18. *p* = 0.0095). **(M)** Distance swam by WT and *gfi1ab^-/-^* larvae with 4 mm gratings at 100% contrast difference (number of larvae: WT n = 17 fish, n *gfi1ab^-/-^* = 18 fish per genotype and condition).

The hunting sequence comprises multiple stages from initiation to the capture of the prey, as previously described in other studies (Fig. 4A) (*39*, *46*, *47*). To further investigate the defects observed in the hunting sequence of DM larvae (Fig. 4B), we recorded 12-minutes videos where single larvae in presence of rotifers performed several hunting sequences (Movies S1, S2). Both WT and DM larvae spent a similar percentage of time foraging (Supplementary Fig. 6C) and hunting with converged eyes (Fig. 4D). Furthermore, there were no significant differences in the initiation of the hunting sequence (Fig. 4E), tracking (Supplementary Fig. 6CI), target fixation (Fig. 4F), or any other steps leading up to predation (Supplementary Fig. 6CII-CC^III^). However, DM larvae showed a significant impairment in initiating capture swims when the prey was within the strike zone (Fig. 4G). Although the mutants were able to approach the prey and get close to it, they often failed to initiate the stereotypical capture swims required for successful predation, opting instead to abort the hunting sequence and abandon the prey.

Additionally, OMR assays indicated that both WT and DM larvae swam similar distances towards high-contrast and bright black and white bars (Fig. 4H), with comparable speeds and initiation latencies (Supplementary Fig. 6D-D^I^). In contrast, mutants swam significantly less towards low-contrast bars compared to WT controls and at a lower speed, highlighting a specific deficit in responding to reduced brightness and/or visual contrast (Fig. 4I, Supplementary Fig. 6D-D^I^). Instead, reducing the size of the moving bars did not affect the OMR response of the DM larvae (Fig. 4J) indicating no defect in visual acuity. The same specific OMR defect was also observed in *gfi1ab^-/-^* mutants (Fig. 4K-M).

These findings demonstrate that the Lrrn2 and Lrrn3a pathway downstream of Gfi1ab plays a critical role in specific visually guided behaviors, particularly those requiring the visual detection of brightness and contrast.

## Discussion

We established Lrrn2 and Lrrn3a as key CAMs regulated by Gfi1ab guiding RGC axons to the SAC/SPV layer of OT, a previously uncharacterized microcircuit that we find essential for brightness and contrast processing. Our findings are particularly significant given that alterations in this molecular pathway impair features of visual behaviors and sensory perception to which we link a particular circuit identified decades ago in vertebrates (*35*).

Lrrn CAMs are evolutionarily conserved both in spatial distribution and functions across species, underscoring their fundamental role in visual system development. In zebrafish and mouse (*48*, *49*), Lrrns are expressed in the retina, while in *Drosophila*, their homologs localize to photoreceptors (*5*, *28*), suggesting shared molecular pathways across vertebrate and invertebrate visual systems (*12*). Functionally, we showed that DM larvae for Lrrn2 and Lrrn3a exhibit highly specific defects in RGC connectivity. While projections to superficial OT layers remain intact, SAC/SPV innervation is specifically disrupted. Moreover, our transplantation experiments reveal that, like Caps, Lrrns operate both cell-autonomously in RGCs and non-cell-autonomously from the OT, ensuring precise target selection. While these findings establish Lrrns as necessary for connectivity in both tissues, we hypothesize that their role likely extends beyond direct homophilic or heterophilic interactions, as previously suggested for Caps (*5*, *17*, *50*). Although CAM overexpression increases aggregation, Lrrns may require *in vivo* additional heterophilic factors to promote or stabilize membrane complexes. This interplay and unique combination of factors likely refines target specificity within microcircuits that express the same molecules (*17*). The partial co-localization of Lrrn expression in the retina, specifically Lrrn2, Lrrn3a, and Lrrn3b, suggests that these proteins may either act redundantly in different combinations or have specialized roles in guiding distinct subsets of RGCs to specific tectal layers. This combinatorial organization resembles the interplay between Caps and its paralog Tartan in *Drosophila*, both contributing to photoreceptor targeting with overlapping but distinct functions (*51*, *52*).

Importantly, the absence of detectable targeting defects in the SAC/SPV layer of *lrrn3b* mutants indicates that its role may be either redundant with other Lrrns or confined to distinct RGC subtypes.

Similarly to Sens controlling Caps in *Drosophila* (*5*, *28*), our analysis identifies Gfi1ab as a key regulator of *lrrn2* and *lrrn3a* in zebrafish, as they partially co-localize with *gfi1ab*-expressing RGCs and are downregulated in the absence of Gfi1ab. When we examined *gfi1ab* mutants, we observed targeting defects within the SAC/SPV layer, providing direct evidence that its regulatory role influences RGC connectivity.

One of the main challenges in studying the SAC/SPV layer has been efficiently labeling this deep region of the tectal neuropile, as transgenic lines like *isl2b*, which mark *pan-RGCs*, do not adequately label the RGCs that project to this layer. Thus, prior functional imaging studies have struggled to directly capture the activity of the SAC/SPV layer, restricting our understanding of its role in higher-order visual processing. Previous efforts to address this gap have involved measuring functional responses from RGCs projecting to intermediate targets, such as AF9 (*27*), or from PVN neurons that project within the deep neuropile (*26*). These studies have suggested a potential role for the SAC/SPV layer in responding to light intensity changes. In our study, the novel *Tg(gfi1ab:gal4;UAS:GcaMP6s)* transgenic line enables us to directly capture RGC activity within the SAC/SPV lamina. We show that this layer is particularly responsive to luminance changes, with higher sensitivity to brightness decreases, especially during sudden or gradual shifts to dark backgrounds (stimuli OFF or dark ramp). Although these neurons respond more strongly to light decrements, they also activate upon presentation of light increment stimuli, suggesting they do not conform strictly to ON or OFF selectivity. These results provide direct evidence that the SAC/SPV layer plays a specialized role in processing luminance shifts, which is essential for detecting dynamic visual cues in low-light environments.

The functional significance of the SAC/SPV layer is further supported by behavioral findings in our DM mutants, which show a higher rate of prey-capture sequence failures, particularly during the critical transition to capture swims when prey is within the strike zone. Although these larvae can detect and approach prey, they fail to execute the precise capture swims necessary for successful hunting. Furthermore, while DM mutants respond robustly to standard and visually demanding OMR stimuli, such as detecting small black-and-white bars, they struggle when contrast is reduced, indicating difficulty in perceiving decreased luminance variations. Recent studies underscore the critical role of brightness and contrast as fundamental optical properties in prey detection, particularly for estimating target distance during the final strike decision (*3*). Our findings align with these observations, as DM larvae, which show defects in prey capture, also exhibit impaired responses to grating stimuli with reduced intensity and contrast. While their ability to detect prey at a distance remains intact, they may fail to accurately estimate its distance as it approaches or struggle to detect it altogether, leading to aborted strike attempts. This suggests that the SAC/SPV layer, which is key in processing luminance cues, might be necessary for distance estimation. Since Lrrn molecules are also expressed in the OT and other brain regions, we sought to confirm whether the observed behavioral deficits stem specifically from retinal neurons. To address this, we tested *gfi1ab^-/-^*mutants, as Gfi1ab expression is restricted to RGCs in the retina and absent in the OT. These mutants exhibited the same prey-capture and OMR deficits as DM larvae, confirming that the observed phenotypes arise from RGC dysfunction rather than Lrrn expression in other brain regions.

The behavioral deficits observed in DM larvae closely resemble those seen when the inter-tectal commissural neurons (ITN) cell bodies are ablated (*39*), hinting at a potential functional link between SAC/SPV-targeting RGCs and the ITNs. These neurons have been shown to be involved in the detection of the prey when located within the binocular strike zone, by enabling a disinhibition circuit necessary for initiating capture swims (*39*). Our findings place ITNs downstream of the SAC/SPV circuit, implying that SAC/SPV-targeting RGCs may indirectly modulate ITN activity. Our connectomics analysis revealed that SAC/SPV-targeting RGCs primarily targeted non-stratified periventricular interneurons (nsPVINs) and projection neurons (nsPVPNs), suggesting downstream connectivity convergence between the SAC/SPV-targeting RGCs and ITNs networks.

Overall, our study unravels a novel neural circuit from genetic and developmental components to functional output and anatomical connectivity, establishing the SAC/SPV layer as a key integrator of luminance processing and distance estimation. By bridging circuit development with visually guided behaviors, we provide new insights into the molecular and structural mechanisms shaping deep tectal connectivity in a vertebrate brain.

## Acknowledgments

We thank all members of the Del Bene lab for discussions and the Institut de la Vision and Institute Curie animal and imaging facilities. We thank Coralie Fassier (Institut de la Vision) for fruitful discussions and for gifting us the *pCX* plasmids, and Stephane Fouquet (Institut de la Vision) for constant help and advice in confocal imaging and IMARIS software. We thank Noé Testa for the help in the initial screening of *gfi1ab*. AI-assisted technology (ChatGPT) was used to correct grammatical mistakes and improve language.

## Author contributions

Conceptualization: EP, FDB, SA Investigation: EP, SA, JBW, FDB Methodology: EP, SA, GF, JTD, FEg, JHS, KD, JV, JBW Visualization: EP, JBW, AB, FEn, JL, SA Funding acquisition: EP, FDB, SA Supervision: FDB, SA Writing – original draft: EP, FDB, SA Writing – review & editing: EP, GF, FEg, KD, JBW, JHS, FDB, SA

## Data and materials availability

All data are available in the main text or the supplementary materials.

## Supplementary Materials

### Materials and Methods

#### Fish Lines and Husbandry

Zebrafish (*Danio rerio*) were maintained under controlled conditions at a temperature of 28°C and subjected to a 14-hour light/10-hour dark cycle with sunset and sunrise. Embryos were collected and cultured in fish water supplemented with 0.003% 1-phenyl-2-thiourea to prevent pigmentation and 0.01% methylene blue (VWR cat n°1.59270.0100) to inhibit fungal growth in the petri dish. The fish were housed within our institute’s animal facility, constructed following local animal welfare standards. All animal procedures adhered to the ethical guidelines outlined by French and European Union regulations. Animal handling and experimental procedures were approved by the committee on ethics of animal experimentation of Sorbonne Université (APAFIS#21323-2019062416186982). Experimental procedures were conducted on larval zebrafish before the onset of sexual differentiation.

#### Zebrafish mutants and transgenic lines

The gRNAs used to generate the different CRISPR knock-in and knock-out lines are listed in **Table S1**, primers used for genotyping are listed in **Table S2**. *Gfi1ab*-/-mutant line was generated using the CRISPR/Cas9 gene editing technology. A gRNA was used to target the first exon of the *gfi1ab* gene leading to a 4 bp deletion, causing a frame shift in the open-reading frame and a premature stop codon at the 6th amino acid. For *lrrn2*-/-mutant line generation, two gRNAs were injected to target the first and single exon of the *lrrn2* gene leading to a large deletion of 323 base pairs, causing the deletion of the signal peptide of the gene. For *lrrn3a*-/-mutant line generation, two gRNAs were injected to target the first and single exon of the *lrrn3a* gene leading to a large deletion of 323 base pairs, causing the deletion of the signal peptide of the gene. For *lrrn3b*^-/-^ three guides were injected to generate a full *knock out* in the F0 generation (*53*), creating a 1000 bp cut and the deletion of most of the LRR domain. PCR amplicons were deposited on 1-3% agarose (LifeTechnologies, cat n° 16500500) gel for the identification of wild type and mutants.

The *gfi1ab:gal4* bacterial artificial chromosome (BAC) construct was generated as follows: a BAC spanning the *gfi1ab* genomic locus (CH211-164G7, BACPAC Resources Center) was used to perform Tol2 transposon-mediated BAC transgenesis replacing *gal4* with the *gfi1ab* coding sequence. Transformation through electroporation of the *pRedET* plasmid was performed as described (*52*). For Tol2 transposon-mediated BAC transgenesis, the iTol2-amp cassette was amplified by PCR with the primer pair pTarBAC_HA1_iTol2_fw (5’- gcgtaagcggggcacatttcattacctctttctccgcacccgacatagatCCCTGCTCGAGCCGGGCCCAAGTG- 3’) and pTarBAC_HA1_iTol2_rev (5’-gcggggcatgactattggcgcgccggatcgatccttaattaagtctactaATTATGATCCTCTAGATCAGCTC- 3’), where the lower and upper cases indicate the *pTarBAC2.1* sequences for homologous recombination and the *iTol2-amp* annealing sequences, respectively. Subsequently, the amplified *iTol2-amp* cassette was introduced into the backbone (*pTarBAC2.1*) of the Gfi1ab-BAC. 500 ng of the PCR product (1 mL) were used for electroporation. 5’- gactgtccgattctcactaatgactggggacatatgaggcagaggaggacGCCACCATGAAGCTACTGTCTTCT ATCGAAC-3’ and 5’ ggctggtggtagctgtgcgcttttttgctcttcaccaagaaagacctaggCCGCGTGTAGGCTGGAGCTGCTTC- 3’ primers were used to amplify and insert the *gal4FF* cassette into the BAC (*55*). The lower and upper cases indicate the CH211-164G7 sequences for homologous recombination and the *pGal4FF-FRT-Kan-FRT* annealing sequences, respectively. 500 ng of the PCR product (1 mL) were used for electroporation in *Gfi1ab-iTol2-amp-BAC*-containing cells. The BAC DNA preparation was performed using the HiPure Midiprep kit (Invitrogen, cat n° K2100), with modifications for BAC DNA isolation as described by the manufacturer. Tol2 transposase mRNA was prepared by *in vitro* transcription from NotI-*linearized pcs2-tol2* (*56*) using the T7 mMessage mMachine kit (Ambion, cat n°Am1348). RNA was purified using the RNeasy purification kit (Qiagen, cat n° 74104), diluted to a final concentration of 100 ng/μl for injection. At least 50 injected fish were backcrossed to wild type. Germline transmission was observed in the offspring from two of such crossings with identical patterns reminiscent of endogenous *gfi1ab* expression.

Along with the newly generated and validated *Tg*(*gfi1ab:gal4*) line, two other transgenic lines expressing *GFP* or *RFP* were used in this study: the previously published *Tg*(*isl2b:GFP*) (*57*); and the previously published *Tg*(*UAS:RFP;cry:GFP*) transgenic line (*58*). The *UAS:GCaMP6s* transgenic line was generated by injecting the *14UASubc:GCaMP6s* plasmid with *tol2* transposase mRNA at 25 ng/µL in one-cell stage zebrafish embryos. To generate the plasmid, we cloned GCaMP6s from plasmid 59530 (Addgene) and inserted it into the 14UASubc backbone, using the NEBuilder® HiFi DNA Assembly Cloning Kit (New England Biolabs, cat n° E5520S) following manufacturer’s instructions.

#### Molecular cloning

The p*UAS:lyn-GFP* plasmid was generated as previously described (*44*). The *pUAS:lynRFP* construct used in this study was previously cloned by Auer and Del Bene (2014) (*56*). The *pDest-Tol2-Isl2b:GFP:polyA* was obtained from Addgene (plasmid n. 105648). The *pCX:lrrn2-eGFP*, *pCX:lrrn3a-RFP* were generated by modifying the *pCX:eGFP* and *pCX:RFP* plasmids from Coralie Fassier. *lrrn2* and *lrrn3a* cDNA was obtained from 3 dpf larvae total RNA extraction. *Lrrn2* and *lrrn3a* cDNA were subsequently subcloned into the *pCX:eGFP* or *pCX:RFP* respectively using the NEBuilder® HiFi DNA Assembly Cloning Kit (New England Biolabs, E5520S) following manufacturer’s instructions. The *pfoxp2.a:Gal4FF* plasmid was obtained from Nikolau and Meyer (2015) (*9*).

#### Micro-injections

Fertilized eggs were collected in egg medium and aligned into prepared 2% agarose (LifeTechnologies, cat n° 16500500) injection molds (agarose diluted in water) (Adaptive Science Tools, cat n°TU-1). To inject, a pressure injector (PicoPump w/pressure (WPI, cat n°PV820), Manual Micromanipulator (WPI, cat n°M3301R) and Injection Assembly parts kit (WPI, cat n°MMP-KIT) with borosilicate glass capillaries (WPI cat n°GBF100-50-10) was used. The capillaries were previously filled with the following injection solutions and injected through the chorion into the cytoplasm of the single cell. The *10UAS:lynRFP* and *UAS:lynGFP* plasmid was injected at 10 ng/μl with or without 20 ng/μl of *tol2* mRNA, either for broad or sparse cellular labelling. The *isl2b:GFP* plasmid was injected at 25 ng/μl with 20 ng/μl of *tol2* mRNA. The *foxp2:gal4FF* plasmid was injected at 15 ng/μl with 10 ng/μl of *tol2* mRNA.

#### Whole mount NBT/BCIP single *in situ* hybridizations

Total zebrafish cDNA ranging from stages 48 hpf to 72 hpf was amplified for probe synthesis. In vitro transcription of Digoxigenin/Fluorescent-labeled probes was carried out using an RNA Labeling Kit (DIG RNA Labeling Mix, cat n° 11277073910; Fluorescein RNA Labeling Mix, cat n°11685619910, Roche) following the manufacturer’s protocols. Zebrafish whole-mount in situ hybridizations were conducted as previously outlined. Subsequently, stained embryos were captured using a stereomicroscope (Leica cat n° MZ10F). Image processing and analysis were conducted using ImageJ software, with uniform adjustments made for color balance, brightness, and contrast.

#### Vibratome Sections

Whole-mount embryos were washed a few times in 1x *PBS*/0.1% *Tween*-20 (PBS-Tw) solution after whole mount NBT/BCIP *in situ* hybridization (Roche, cat n° 11681451001). The samples were embedded in gelatin (Sigma Aldrich cat n° G1890)/ albumin (Sigma Aldrich, cat n° A4503) with 4% of Glutaraldehyde (Sigma Aldrich, cat n° 340855) and sectioned (20 mm) on a vibrating blade microtome (Leica, cat n° VT1000 S). Sections were mounted in Fluoromount Aqueous Mounting Medium (Sigma Aldrich, cat n°F4680).

#### TUNEL Assay

To detect apoptosis on whole mount zebrafish larvae at 4, 5 and 7 dpf the TUNEL assay was employed, using Apoptag Peroxidase In Situ Apoptosis Detection Kit (Millipore, cat n° S7100). Larvae at 4, 5 and 7 dpf were fixed in 4% PFA (Electron Microscopy Sciences, cat n° EM-15710) diluted in PBS (Euromedex cat n° EU1-9400-100) with 0.1% Tween-20 (Sigma-Aldrich, cat n° 0777) for 2h at RT and then dehydrated and stored in methanol (Fisher Scientific, cat n° 15623710) at –20°C. Dehydrated larvae fixed at the proper developmental stages were rehydrated at RT using successive and decreasing dilutions of methanol in PBST. The samples were then washed in PBST. To allow the permeabilization of the samples and the penetration of the probe the larvae were digested at RT with 10 μg/mL of Proteinase K (Sigma-Aldrich, cat n° 03115852001) diluted in PBST for 45 or 60 minutes, according to their developmental stage.

Proteinase K digestion was stopped by incubating the samples in 4% PFA (in PBST) for 20 minutes or more at RT. Several washes were carried out using PBST before incubating the larvae in 100%EtOH (VWR cat n° 84857-360)/100%Acetic Acid (VWR, cat n°20104.298), mixed 2:1, at-20°C for 15 minutes. Several washes were then performed in PBST at RT. The larvae were then equilibrated in the Equilibration Buffer provided by the kit at RT for 15 minutes. The samples were then transferred to terminal transferase enzyme solution mixed with Reaction Buffer 1:2 (both in the Apoptag kit), and 3% TritonX-100 (Euromedex cat n° 2000-C)/PBS for 1 hour on ice and then for 1 hour at 37°C. 1% Stop Solution (from the kit) was used to stop the transferase reaction for 5 minutes at RT and then for 45 minutes at 37°C. Several washes were then carried out in PBST. The larvae were then incubated for at least 1 hour in 10% Blocking Solution (Roche, cat n° 11096176001) in order to saturate the non-specific binding sites before adding antibodies anti-DIG-Fab-Fragments conjugated to alkaline phosphatase (Roche, cat n° 11093274910) diluted 1/5000 in 2x Blocking Solution. The samples were incubated in this solution overnight at 4°C. A series of PBST washes were performed the next day before incubating the samples in the Staining Buffer (5M NaCL, 1M MgCl2, 1M Tris-HCl (pH 9.5), 100% Tween-20, H2O) twice for 10 minutes at RT. The color reaction was then carried out by incubating the larvae with NBT/BCIP (Roche, cat n° 11681451001) diluted 1/500 in the Staining Buffer in total darkness for as long as necessary. To stop the reaction the larvae were washed several times with PBST, and then fixed overnight at 4°C in 4% PFA in PBST. The samples were then analyzed on ZEISS SteREO Discovery.V20 microscope after an overnight incubation at 4°C in 87% glycerol (VWR cat n° 24388.295) diluted in water.

#### Cryo-sectioning

Zebrafish larvae at the developmental stage of 3 or 5 dpf were fixed in 4% paraformaldehyde in 1x phosphate-buffered saline (PBS; pH 7.4) for 2 hours at room temperature and, thereafter, cryo-protected overnight in a 30% sucrose (VWR, cat n° 27480.294)/0.02% sodium azide (Sigma, cat n° S2002) /PBS solution. Embryos were transferred to plastic molds and embedded in O.C.T. Compound (Euromedex, cat n° 62550-12) after removal of the sucrose. Blocks were then frozen at-80°C on dry ice. The 14 μm sections were mounted on Superfrost Plus slides (Fisher Scientific, cat n° 10149870).

#### Immunohistochemistry

For Anti-GFP immunohistochemistry no antigen retrieval was performed and 1/500 dilution of chicken primary anti-GFP antibody (Genetex, cat n° GTX13970) and Alexa Fluor 488 secondary antibody goat anti-chicken IgG (1/1000, LifeTechnologies, cat n° A-11041) were used.

Immunohistochemistry experiments were performed as follows: cryosections were washed three times in 1×3 PBS/0.1% Tween-20 (PBS-Tw) solution. Consecutively, slides were incubated 1 hour at room temperature in 10% blocking solution (Roche) followed by overnight incubation at 4°C in 1% blocking solution (10% blocking solution in 1xPBS-Tw) in which primary antibodies were diluted. Sections were then washed five times using 1×3PBS-Tw solution and incubated at room temperature for 1 to 2 hours in 1% blocking solution in which primary antibodies were diluted. Slides were then washed several times in 1xPBS-Tw solution, counterstained with DAPI (LifeTechnologies, cat n° D3571) diluted in 1xPBS-Tw solution for 10 minutes. The DAPI solution was then washed out several times in 1xPBS-Tw solution. Slides were then mounted using an 80% glycerol solution prior to imaging.

#### HCR

All HCR probes and buffers were purchased from Molecular Instruments®. 48 – 72 hpf larvae underwent fixation in 4% paraformaldehyde in PBS (pH 7.4) for 2 hours at room temperature, followed by multiple rinses with PBS to halt the fixation process. Subsequently, larvae were subjected to dehydration through a series of methanol (MeOH) washes and then stored at-20°C. The HCR procedure was carried out according to the manufacturer’s instructions. HCR fluorescent samples were then imaged using a 40x inverted immersion objective (NA 1), with Z-volumes captured at a resolution of 1.5 μm, using an Olympus FV3000 confocal microscope.

Image processing and analysis were conducted using ImageJ and Cellpose software. The first was used to extract maximum projections of 2 to 4 stacks while the latter to segment DAPI signal in order to retrieve nuclei’s outline membrane borders. These membrane outlines were used to better identify cells co-expressing RNAs of interest.

#### Transplantation

Mutant and wild-type chimeric embryos were obtained by blastomere transplantations at the 1000-cell stage as previously described (*57*). Donor RGCs were derived from *Tg(lrrn2^-/-^;lrrn3a^-/-^*;*gfi1ab:gal4)* mutant or *Tg(lrrn2^+|+-^;lrrn3^+/+^;gfi1ab:gal4)* wild-type embryos, previously injected at 1 cell stage with *UAS;lyntagRFP* and transferred into either *Tg(isl2b:GFP)* or *Tg(lrrn2^-/-^;lrrn3a^-/-^;gfi1ab:gal4;isl2b:GFP)* acceptor embryos of the same stage.

#### Aggregation Assay

The Aggregation Assay was performed as described (*60*). To evaluate homophilic interaction, HEK293 cells were independently transfected with *pCX:eGFP, pCX:mRFP, pCX:lrrn2-eGFP, and pCX:lrrn3a-mRFP*. Images of three random areas per well were acquired for each sample using a Leica DM6000 Epifluorescent Microscope with a 10X objective. Transfected and resuspended cells expressing each construct, not subjected to aggregation, were transferred to a 6-well plate and used as controls.

Fluorescent signals from single cells or aggregates were detected and counted semi-automatically using a custom-made MATLAB code. This code detected and highlighted structures based on a pixel intensity threshold. The detected structures were circled with colors that varied based on the radius of each structure. Various images were also manually evaluated to validate the code. Images of the three random areas per well were averaged together. The aggregation index was calculated by dividing the number of aggregates present in the control wells by the number of aggregates present in the aggregated wells.

To evaluate heterophilic interaction, cells were independently transfected with *pCX:eGFP, pCX:mRFP, pCX:lrrn2-eGFP, and pCX:lrrn3a-mRFP*. Before starting the aggregation protocol, *pCX:eGFP*-transfected cells were mixed with *pCX:mRFP* cells, and *pCX:lrrn2-eGFP*-transfected cells were mixed with *pCX:lrrn3a-mRFP* cells and imaged as controls. Image acquisition was performed as before, and homophilic interaction was tested as well. To evaluate possible interaction between Lrrn2 and Lrrn3a, we manually counted the aggregates expressing both red and green fluorophores and normalized the total number of co-expressing aggregates after aggregation to the number of co-expressing aggregates before aggregation.

#### RNA extraction and cDNA preparation and qRTPCR

Total RNA extraction was conducted from dissected heads of 2 and 3 dpf zebrafish larvae using TRIzol reagent (LifeTechnologies, cat n° 15596018) and TURBO DNA-free reagents (Ambion AM1907). The RNA was derived from a mix of the heads of 50 larvae of the same genotype.

The concentration of RNA was determined using the Nanodrop2000 (Thermo Fisher). For reverse transcription, 1 μg of RNA was retro-transcribed using random primers and the SuperScript III First-Strand Synthesis system (LifeTechnologiesn cat n°18080051). SYBR Green PCR Master Mix (Ambion, cat n° 4368702) was employed for qRT-PCR on QuantStudio 6 PCR System instrument (ThermoFisher) according to the manufacturer’s instructions. *Ef1a* and *rpl13a* were utilized as reference genes, as previously documented. All assays were conducted in technical triplicates and replicated in three independent biological experiments. Mean values from triplicate experiments were determined using the delta CT quantification method, with p-values calculated using a Mann-Whitney non parametric test. The primers used for qRTPCRs are in **Table S3**.

#### Comparison between *Tg(gfi1ab)* and *Tg(isl2b)* lines

Fluorescence intensity was quantified using ImageJ (NIH) to compare visualization of the SAC/SPV layer in *Tg(gfi1ab)* and *Tg(isl2b)* transgenic fish lines. For each image, obtained through confocal microscopy, pixel intensity was measured in both the SFGS and SAC/SPV layers. To normalize fluorescence across images and reduce image-to-image errors, a fluorescence index was calculated by dividing the mean intensity of SAC/SPV by the mean intensity of SFGS within each image. This analysis was performed in both transgenic lines to assess differences in imaging quality.

#### Imaging of sparse RGCs and PVNs

Mutant and WT control RGC axons were acquired between 3 and 5 dpf using a 40X Wplan Apochromat objective (NA 1; WD 2.5mm), with Z-volumes captured at a step size of 1.5 μm, using an Olympus FV3000 laser scanning confocal microscope. Image acquisition was conducted at a resolution of 1024×1024 pixels, with a scan rate of 4 μs/pixel. The same imaging approach was used for the transplantation experiments.

#### Single RGC Analysis

To analyze signal from the *Tg(gfi1ab:gal4)* transgenic line the 3D reconstructions of the Z-volumes of RGCs axons were obtained using the Oxford Imaris Suite software 10.0. 3D images were rotated in order to visualize the synaptic laminae. Skin autofluorescence and the signal from the retina were removed manually. To isolate single RGCs axons, we utilized the surface tool with semi-automatic selection and then applied the surface to mask the RGC when its projections were clearly distinguishable from other axons. We then used the filament tracing tool to properly trace the RGC and calculate its absolute distance from a reference point positioned manually. Such reference point was consistently positioned on the skin and 3D oriented using the signal from the *Tg(isl2b:GFP)* background. When traced RGCs presented several arborizations, we calculated independently the absolute distance from the tip of each arborization to the reference point, which were then averaged to obtain a unique value for each RGC. Depending on the distance from the skin, each RGC was identified as SO, SFGS or deep-projecting RGC. Statistical analysis was conducted separately for RGCs projecting to different layers. The percentage of correct and mistargeted deep-projecting RGCs was also assessed and compared between mutants and WT controls.

To analyze the projections of RGCs labelled by the *Tg(isl2b:GFP)* plasmid, we used syGlass, a virtual reality software, in combination with an Oculus Quest (Meta) headset. This approach was essential, as conventional methods using Imaris alone were insufficient for the detailed tracking of individual axon projections. In syGlass, we used the Region of Interest (ROI) tool to manually track and segment the axons in 3D and generate masks. The final segmented images were saved as TIFF files, which were compatible with Imaris 10.0, allowing us to visualize and manipulate multiple channels/masks. The datasets were organized to maintain the same dimensionality as the original data, enabling precise visualization of the axon projections with distinct pseudo-color assignments.

#### Single PVN Analysis

The 3D reconstructions of the Z-volumes of PVNs were obtained using the Oxford Imaris Suite software. 3D images were rotated in order to visualize the neuron’s morphology. Skin autofluorescence and the signal from the retina were removed manually. The quantification of stratified vs non-stratified neurons was performed as in Di Donato et al., 2018 (*42*).

#### Behavioral tests

- free swimming

Free swimming behavioral acquisitions, fish tracking, tail segmentation and analysis were performed as previously described (*61*). In total 12 larvae were tested for each genotype with the acquisition lasting for 20 to 30 minutes.

- OMR

To assess the optomotor response (OMR) in our fish, we employed the Stytra software platform

(*62*). The fish were tracked using Stytra, which provided precise tracking data, and the videos were recorded at 100 Hz. The same camera and illumination setup as in the free-swimming experiment was used. Visual stimuli were projected below the 60 mm petri dish via a cold mirror. The visual stimuli consisted of sinusoidal gratings of 4 and 10 mm spatial frequency and a speed of 10mm/s. High contrast bars were achieved by presenting black and white gratings, set as 100% of difference of contrast intensity, while low contrast stimuli by decreasing the contrast to 20% of difference. The stimulus consisted of 6 repetitions of 20 seconds of either left-ward and right-ward movements, alternating after 20 second pauses. The larvae were left to adapt to a light illuminated background before stimulus presentation for over 15 minutes. Subsequently, we utilized a custom-made Python code to extract detailed parameters from the tracking data, including trajectories, speed, tail beat frequencies, distances, and latency.

- Prey capture assay

Prey consumption assays were conducted at 6 days post-fertilization and performed as previously described (*37*).

- Prey capture sequence

To assess the outcomes of individual hunting sequences, 12-minute videos capturing hunting larvae were recorded at a rate of 100 Hz. Dark-field illumination was achieved by using a custom-made LED-ring (wavelength = 850 nm) positioned around the dish, facilitating the detection of rotifers. Hunting sequences were identified based on observable eye vergences and categorized according to their outcomes.

#### Zebrafish larvae used for two-photon and EM connectomic atlas

The triple transgenic *Tg(elavl3:H2B-GCaMP7f^+/+^, gad1b:dsRed^+/-^, flk1:mCherryCAAX^+/-^)* larvae used in this study was obtained by crossing adult transgenic *Tg(elavl3:H2B-GCaMP7f, gad1b:DsRed)* and *Tg(elavl3:H2B-GCaMP7f, flk1:mCherryCAAX)* lines in nacre background, *mitfa^−/−^*(*63*) yielding high GCaMP7 fluorescence. The functional connectomics experiment was performed on 7 dpf larvae. For single-cell electroporation we incrossed *Tg(elavl3:H2B-GCaMP7f)* lines. All experiments were approved by the Harvard University standing committee on the use of animals in research and training. Larvae were fully embedded in a drop of agarose (UltraPure Low Melting Point Agarose, Invitrogen cat n° 16520050), at ∼35 °C at the center of a Petri dish (9 cm in diameter, Dutscher cat n° 633180). After the agarose was solidified the dish was filled with fish facility water and transferred into the measurement chamber of a custom-built two-photon microscope, operated by custom-written Python 3.7-based software (PyZebra2P). We used a femtosecond-pulsed laser (MaiTai Ti:Sapphire, Spectra Physics) equipped with a set of x/y-galvanometers (Cambridge Technology), a 20× infrared-optimized objective (XLUMPLFLN, Olympus) to scan over the brain and two photomultipliers (green and red) amplified by two current preamplifiers (SR570, Stanford) to collect fluorophores emissions.

#### Sample preparation for the EM imaging

The following protocol was specifically modified from (*64*) to enhance the extra-cellular space preservation which improves synapses detection and permit single cell resolution imaging using X-ray tomography. Unless noted, all steps were performed at room temperature (RT).

Immediately after the two-photon stack acquisition, the larva still anesthetized and embedded in agarose, we replaced the water by a dissection solution (64 mM NaCl, 2.9 mM KCl, 10 mM HEPES, 10 mM glucose, 164 mM sucrose, 1.2 mM MgCl2, 2.1 mM CaCl2, pH 7.5) supplemented with 0.02% tricaine (Sigma Aldrich, cat n° E10521) (*65*). We then cut small slits in the agarose to expose the eyes and performed bi-lateral enucleations to enhance ultrastructural preservation (extra-cellular space) and heavy metal staining. We used a custom-made hook that was carefully inserted behind the eyes to prevent brain damage. The larva was immediately transferred to a fixation solution at 4°C (2.5% glutaraldehyde, 0.1M cacodylate buffer supplemented with 4.0% mannitol at pH7.4). Cacodylate buffer: 0.3M sodium cacodylate, 6mM CaCl2, pH7.4. To improve the fixation the tissue was rapidly microwaved (cat. no. 36700, Ted Pella, equipped with power controller, steady-temperature water recirculator and cold spot) in the fixative solution (this step lasted <5 min after initial transfer into fixative). The microwaving sequence was performed: at power level 1 (100 W) for 1 min ON, 1 min OFF, 1 min ON then to power level 3 (300 W) and fixed for 20 s ON, 20 s OFF, 20 s ON, three times in a row. Fixation was then continued overnight at 4°C in the same solution. The following day, the sample was then washed again in 0.5x cacodylate buffer (3 exchanges, 30 min each before osmication (2% OsO4 in 0.5x cacodylate buffer, 90 min). After a quick wash (< 1 min) in 0.5x cacodylate buffer the sample was reduced in 2.5% potassium ferrocyanide in 0.5x cacodylate buffer for 90 min then washed with filtered H_2_O (3 exchanges, 30 min each) and then incubated with 1% (w/v) thiocarbohydrazide (TCH) in filtered H_2_O (and filtered with a 0.22 μm syringe filter before use) for 45 min to enhance staining (Hua et al., 2015). Due to poor dissolution of TCH in water, the solution was heated at 60°C for ∼90 min with occasional shaking before filtering and then placed at RT for 5min before the incubation step. The sample was then washed with filtered H_2_O (3 exchanges, 30 min each) before the second osmication (2% OsO4 in filtered H_2_O, 90 min) and then washed again (3 exchanges, 30 min each). Then, en-bloc staining was performed overnight using 1% uranyl acetate in filtered water at 4 °C. The solution was sonicated for 90 min and filtered with a 0.22 μm syringe filter before use. Steps involving uranyl acetate were performed in the dark. The following day, samples were then washed with filtered H_2_O (3 exchanges, 30 min each). Next, the samples were dehydrated in serial dilutions of ethanol (25%, 50%, 75%, 90%, 100%, 100% for 10min each step) then in propylene oxide (PO) (100%, 100%, 30min each step). Infiltration was performed using LX112 epoxy resin with BDMA (21212, Ladd) in serial PO dilutions steps, each lasting 4h (25% resin/75% PO, 50% resin/50% PO, 75% resin/25% PO, 100% resin, 100% resin). Samples were mounted in fresh resin in a mouse brain support tissue with the head exposed to facilitate cutting by avoiding the sample to sink to the bottom of the resin molds. Mouse tissue was fixed using standard procedures (*66*) then cut into 2-3 mm wide cubes which were pierced using a puncher (0.75 mm, 57395, EMS) to insert the larva. The cubes were stained along with fish samples using the protocol described above except that the uranyl acetate overnight step was performed at RT. The samples were then cured with support tissue during 3 days at 60°C. For all steps, a rotator was used. Aqueous solutions were prepared with water passed through a purification system (Arium 611VF, Sartorius Stedim Biotech). The protocol lasted 5 consecutive days including surgery, fixation, staining, and resin embedding followed by 3 days of resin curing.

#### mSEM Image alignment, stitching and rendering

We trained convolutional neural networks to detect defects and classify locations that contain tissue for each montaged section. We used self-supervised convolutional neural networks to generate dense displacement fields between sections (*67*), then hierarchically minimized the elastic energy in the set of all displacement fields to produce an aligned image. We created an “image mask” that contained defects and locations that were classified as misaligned. We trained convolutional neural networks that we used to segment cells, detect synaptic clefts, and assign presynaptic and postsynaptic objects from the cell segmentation at each cleft (*68*). Cell segmentation and synapse detection and assignment used the aligned image with locations in the “image mask” set to black. The cell segmentation was ingested into the ChunkedGraph proofreading system so that errors could be manually corrected, and synapses were ingested into an annotation table for consistent analysis (*69*).

#### Calcium Imaging

To monitor calcium dynamics, we used a commercial 2P scanning microscope (LAVision, Miltenyi Biotec) coupled with a tunable Ti:Sapphire femtosecond laser (Coherent Chameleon, wavelength range 700–1100 nm). Calcium transients were acquired via bidirectional galvo-galvo scanning at 920 nm, with a scanning rate of 3 Hz and a field of view of 300 × 300 μm.

#### Visual stimulation for calcium imaging

Visual stimuli were delivered using a red-emitting projector, projecting on a diffusive screen positioned at ∼3 cm from the larva’s head and covering 120° horizontal visual field. The visual stimuli sequence consisted in:(i) Bright Flash: a single 10 second bright flash stimulus; (ii) Gratings: four gratings corresponding to the cardinal directions (0°, 90°, 180°, and 270°) with 10°-wide dark stripes moving at 20°/s. Each grating was presented statically in the first set of stimuli, then in motion for 5 seconds in the second set of stimuli); (iii) Bright Ramp: a gradual increase in brightness over 5 seconds; (iv) Looming Stimuli: a steady looming, a dark circular dot whose dimension expands at a constant rate of 20°/s, followed by an exponential looming, with lateral dimensions doubling every second); (v) Dark Ramp: a gradual decrease in brightness over 5 seconds. A second sequence consisted in: (i) Prey-dot stimuli: a 5 µm dot moving horizontally four times back and forth at a speed of 80°/sec with the stimulus repeated twice. (ii) Gratings: same as in the first sequence. Stimuli were programmed and synchronized with calcium imaging recordings, ensuring precise timing and reproducibility.

#### Analysis of visually induced calcium responses

Single-ROI traces were extracted using the Suite2p software, applying its functional detection module following motion correction. ROI detection was performed with a threshold scaling factor of 0.75 and a maximum allowed ROI overlap of 1. Raw fluorescence traces (F) were extracted for each ROI, and neuropil correction was applied using the default coefficient (0.7 × Fneu). ROIs located within the same neuropil layer—manually identified based on anatomy and their distance from the skin—were normalized to their respective baseline values (mode of the raw fluorescence trace) before averaging across ROIs to yield a single calcium trace per layer, per fish. Calcium transients were defined as ΔF/F = (F(t) - F₀)/F₀, with F₀ being the mode of the trace. The resulting average calcium transients per layer were further normalized to their respective maximum values. Finally, normalized traces from individual fish were averaged to obtain a single representative calcium transient for each layer. These normalized traces were subsequently analyzed using customized functionalities of the OASIS package within a MATLAB environment (*70*). A first full trace deconvolution was performed based on a second-order autoregressive kernel, constructed as the difference between a rising and a decaying exponential. The resulting neuronal activity trace (a set of discrete peaks representing the individual events that generate the overall calcium transients) was then decomposed into multiple time windows, defined by the delivery of specific visual stimulation to the eye of the fish.

Subsequently, activity peaks corresponding to the stimuli of interest were reconvolved with the kernel function to evaluate the contribution of each stimulus to the global calcium responses, and to separate it from interstimulus crosstalk, as well as from the effects of transitions between stimulation patterns and spontaneous activity. The peak values of each stimulus contribution were then used to quantitatively compare the magnitude of responses across layers and visual stimuli.

#### Statistical Methods

All comparisons between WT controls and mutant larvae were carried out on Prism 7 (Graphpad). To assess statistical significance the Mann-Whitney test by ranks was performed when the dataset did not follow a normal distribution or did not meet basic t-test assumptions.

---

**Supplementary Fig. 1.**
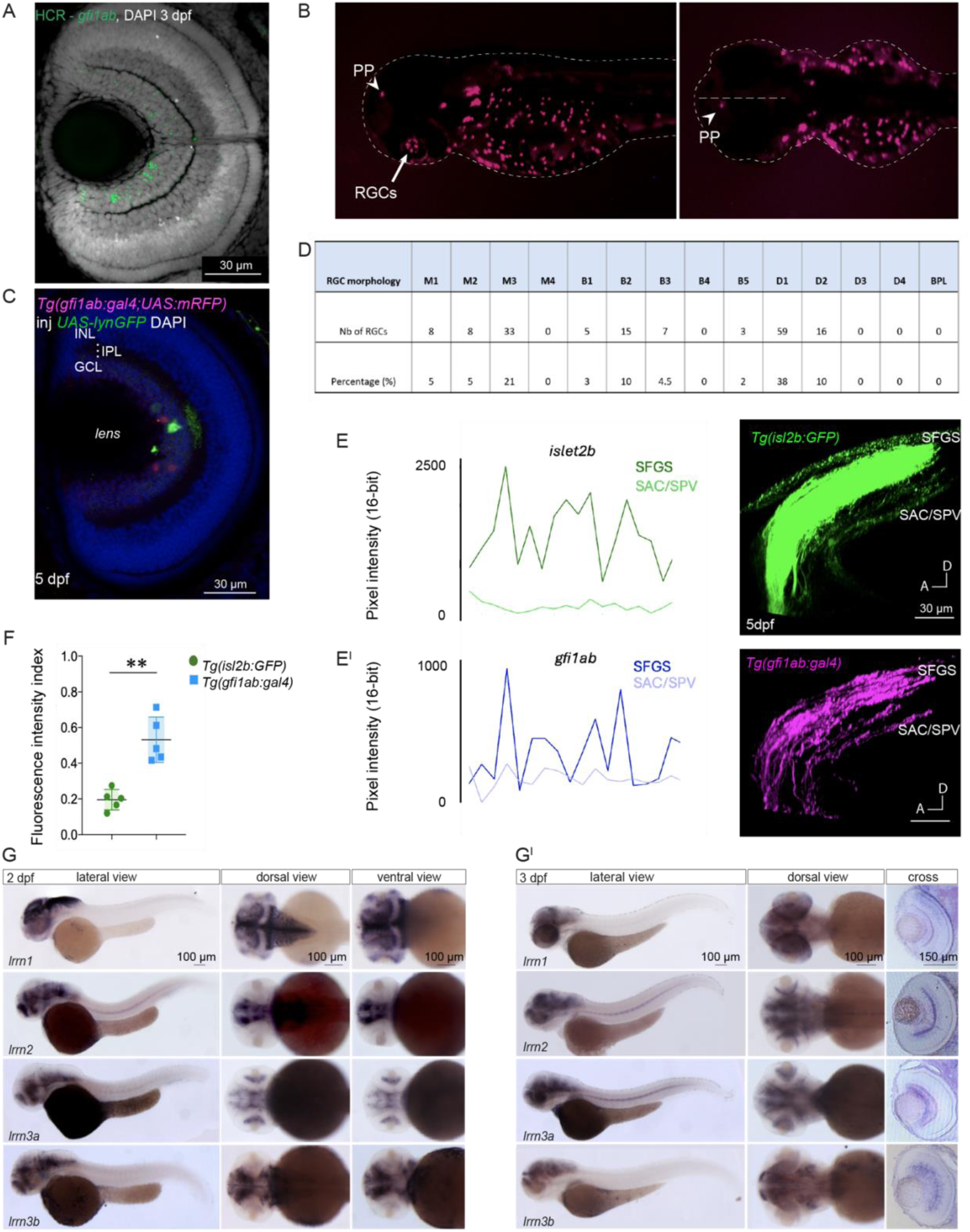
(A) *Gfi1ab* sparse expression in RGCs at 3 dpf. Confocal stack of a 3 dpf zebrafish retina stained by whole-mount HCR *in situ* hybridization for *gfi1ab* (green) and DAPI (grey) highlighting cell nuclei. **(B)** The *Tg(gfi1ab:gal4)* transgenic line recapitulates endogenous *gfi1ab* expression in the RGCs, pineal gland, and ear hair cells. **(C)** Confocal single stack of a 5 dpf zebrafish retina, showing *gfi1ab*-expressing RGCs dendritic stratification in the IPL. **(D)** Quantification of dendritic Gfi1ab-RGC morphotypes in number and percentages identified in 5 dpf *Tg(gfi1ab:gal4;UAS:mRFP)* embryos transiently injected with a *10UAS:lynGFP* plasmid for sparse labeling of single Gfi1ab-RGCs at the membrane (green). Classification follows previously identified morphological groups (*11*). **(E-E^I^)** Comparison of fluorescent intensity index in the *Tg(isl2b:GFP)* line (E) and the *Tg(gfi1ab:gal4;UAS:RFP)* line (E^I^). Pixel indexes were obtained by normalizing the mean fluorescence pixel of the SAC/SPV layer to the mean index of the SFGS within the same fish (n = 5 larvae per transgenic line). **(F)** Dot plot showing fluorescent intensity index of the mean fluorescence intensity of the SAC/SPV layer normalized to the mean fluorescence intensity in the SFGC layer of the *Tg(isl2b:GFP)* and the *Tg(gfi1ab:gal4;UAS:mRFP)* transgenic lines respectively (mean intensity for *Tg(isl2b:GFP) = 0.5,* for *Tg(gfi1ab:gal4;UAS:mRFP = 0.2*). **(G-G^I^)** NBT/BCIP whole mount *in situ* hybridization of all four *lrrn* isoforms at 2 dpf (G) and 3 dpf (G^I^) reveals broad expression in the forebrain, midbrain, and hindbrain. Cross sections of the retina at 3 dpf show differential expression patterns within the neural retina of *lrrn1*, *lrrn2*, *lrrn3a* and *lrrn3b*. Scale bars = 50 μm; Horizontal bars = mean ratios; Vertical Bars = standard deviation. Abbreviations: Tg, transgenic; PP, parapineal gland; RGC, retina ganglion cells; INL, inner nuclear layer; IPL, inner plexiform layer; GCL, ganglion cell layer; Nb, number; inj, injected. Statistical significance was determined in all cases using a non-parametric Mann-Whitney U-test and depicted as: ns – *p* > 0.05; * *p* < 0.05; *\*\* p < 0.01*.

**Supplementary Fig. 2.**
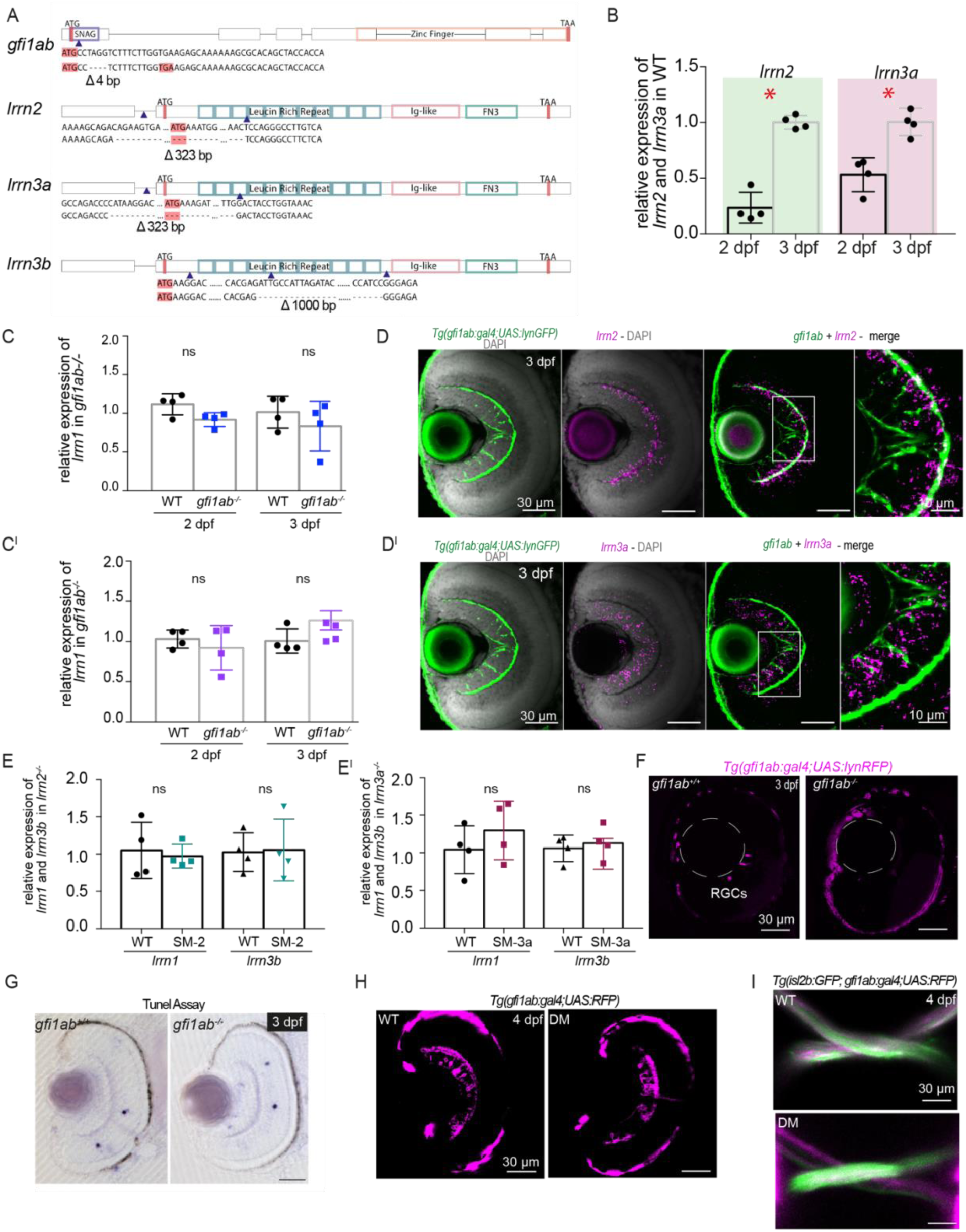
(A) Schematic representation of the deletions and targeted regions encoding for specific protein domains of *gfi1ab*, *lrrn2*, *lrrn3a*, and *lrrn3b* loci. The generated loss-of-function mutations are annotated below each gene (D). Red rectangles highlight the start (ATG) and stop (TAA) codons. Blue triangles indicate the gRNA cut sites. WT sequence around the start codon is compared with the sequence after gRNA-induced mutations. (**B)** Quantitative RT-PCRs performed on the heads of 2 and 3 dpf WT show strong increase in *lrrn2* and *lrrn3a* expression levels from 2 to 3 dpf. n = 3 technical replicates, 50 heads per condition, n = 4 biological replicates per developmental stage and genotype; *p* = 0.0286). (**C-C^I^)** Quantitative RT-PCRs performed on the heads of 2 and 3 dpf *gfi1ab^-/-^* mutants show no difference in *lrrn1* (C) and *lrrn3b* (C^I^) expression levels. n = 3 technical replicates, 50 heads per condition, n = 4 biological replicates per developmental stage and genotype. C. Quantitative RT-PCRs performed on the heads of 3 dpf *lrrn2^-/-^* mutants show no difference in *lrrn1* and *lrrn3b* expression levels. n = 3 technical replicates, 50 heads per condition, n = 4 biological replicates per developmental stage and genotype. (**D-D^I^)** Confocal Z-stacks of 3 dpf *Tg*(*gfi1ab:gal4;10UAS:lynGFP*) larval retinae counterstained for *lrrn2*, and *lrrn3a* HCR probes. *Gfi1ab*-expressing cells are labeled using a *10UAS:lynGFP* plasmid labeling the membranes with GFP (D, D^I^). HCR confirms the expression of *lrrn2* (D) and *lrrn3a* (D^I^) in the GCL. Merged images (D and D^I^) show partial overlap of *lrrn2* and *lrrn3a* signals in *gfi1ab*-expressing GFP cells respectively. Magnifications of double-stained cells. (**E-E^I^)** Quantitative RT-PCRs performed on the heads of 3 dpf *lrrn3a^-/-^*mutants show no difference in *lrrn1* and *lrrn3b* expression levels. n = 3 technical replicates, 50 heads per condition, n = 4 biological replicates per developmental stage and genotype. (**F)** Confocal section of 3 dpf *Tg(gfi1ab:gal4;UAS:mRFP)*;*gfi1ab^+/+^* embryos. RFP-positive RGCs can already be detected at this developmental stage in contrast to *Tg(gfi1ab:gal4;UAS:mRFP);gfi1ab^-^*^/-^ where no mRFP was observed in the retina (n = 6 for both genotypes). (**G)** No difference in cell death could be observed at this stage in *Tg(gfi1ab:gal4;UAS:mRFP)*;*gfi1ab^+/+^*embryos and *Tg(gfi1ab:gal4;UAS:mRFP);gfi1ab^-^*^/-^ embryos as shown by TUNEL assay (n = 8 for both genotypes). (**H)** Confocal imaging of a 3 dpf retina of a WT (G) and DM larva (G^I^), in the *Tg(gfi1ab:gal4;UAS:mRFP)* background line. No gross morphological changes are observed in DM larvae compared to WT controls (n = 7 for WT, n = 5 for DM). (**I)** Confocal imaging of the optic tract of a 4 dpf *Tg(gfi1ab:gal4;UAS:mRFP)* WT (H) and DM larva (H^I^). No gross morphological changes are observed in the DM larvae compared to WT controls (n = 4 for WT, n = 3 for DM). Scale bar = 30 μm. Horizontal bars = mean ratios; Vertical Bars = standard deviation. Abbreviations: WT, wild-type; DM, double mutant; ns, not significant. Statistical significance was determined in all cases using a non-parametric Mann-Whitney U-test and depicted as: ns – *p* > 0.05; * *p* < 0.05.

**Supplementary Fig. 3.**
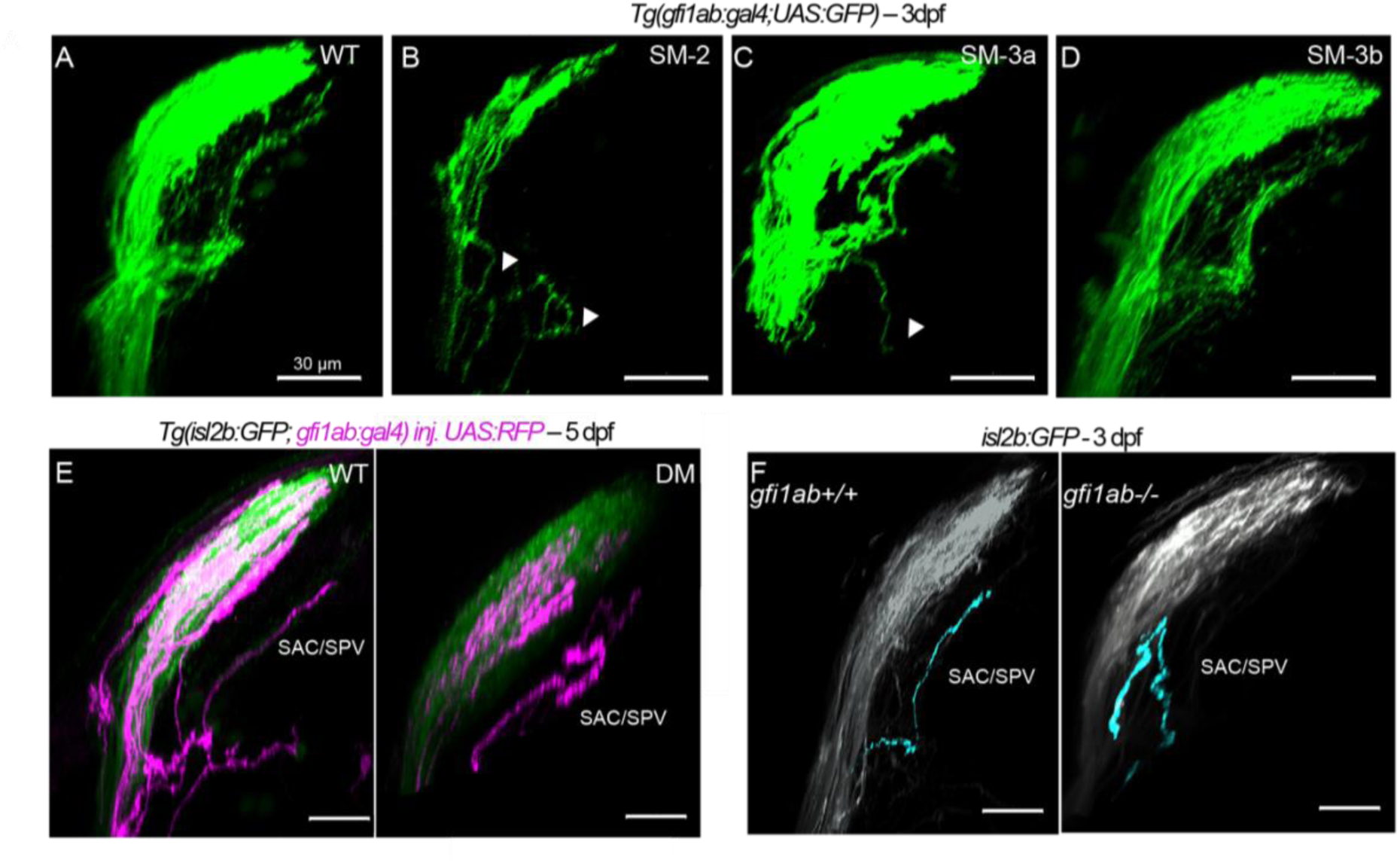
(A-D) Confocal imaging of 3 dpf RGC axonal projections to the optic tectum (OT) in WT (A), SM-2 (B), SM-3a (C), and SM-3b (D)*Tg(gfi1ab:gal4;UAS:GFP)* larvae. Mistargeting defects are detected in SM-2 and SM-3a RGCs axons projecting to the SAC/SPV (mistargeting defects indicated by white arrowheads). In contrast, no mistargeting defects were observed in SM-3b (n = 12 per genotype). **(E**) Confocal imaging of 5 dpf RGCs in a WT (E) and DM (E^I^), in the *Tg(gfi1ab:gal4;UAS:mRFP)* larvae. SAC/SPV mistargeting defects are still detected at this developmental stage (white arrowheads) (n = 15 WT, n = 7 DM). **(F)** Confocal imaging of 3 dpf RGC axonal projection to the OT in WT and *gfi1ab^-/-^*larvae, after injection of *isl2b:GFP* plasmid in 1-cell stage embryos, showing mistargeting defects in the SAC/SPV layer of the mutants. Scale bars = 30 μm. Abbreviations: WT, wild-type, SM, single mutant, DM, double mutant.

**Supplementary Fig. 4.**
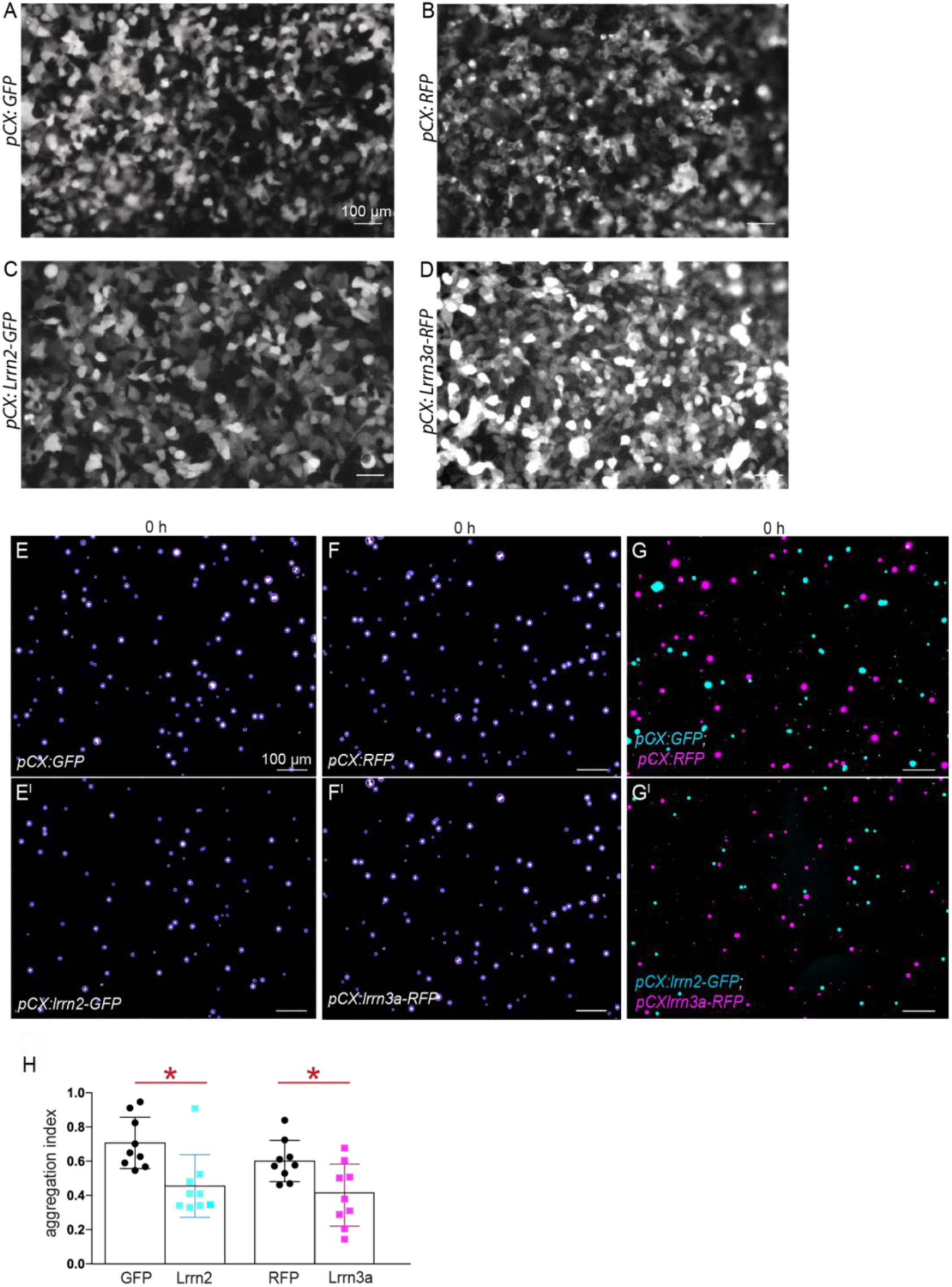
(A-D) Epifluorescence imaging of undissociated HEK293 cells prior to the aggregation assay, transfected with *pCX:GFP* (A), *pCX:RFP* (B), *pCX:lrrn2-GFP* (C), and *pCX:lrrn3a-RFP* (D). **(E-F^I^)** HEK293 cells transfected with *pCX:GFP* (E) and *pCX:RFP* (F), as well as *pCX:lrrn2-GFP* (E^I^) and *pCX:lrrn3a-RFP* (F^I^), imaged at timepoint 0 (0 h). **(G-G^I^**) HEK293 cells mixed after transfection with either *pCX:GFP* and *pCX:RFP* (G) or *pCX:lrrn2-GFP* and *pCX:lrrn3a-RFP* (G^I^), imaged at timepoint 0 before the aggregation assay. **(H)** Dot plot showing the aggregation index of cells transfected and then co-mixed with *pCX:GFP* (cyan) and *pCX:RFP* (magenta) (G) compared to those transfected with *pCX:lrrn2-GFP* and *pCX:lrrn3a-RFP* and mixed together at timepoint 0. Scale bar = 100 μm. Horizontal bars = mean ratios; Vertical Bars = standard deviation. Statistical significance was determined using a non-parametric Mann-Whitney U-test. Data are represented as mean. \**p* < 0.05.

**Supplementary Fig. 5.**
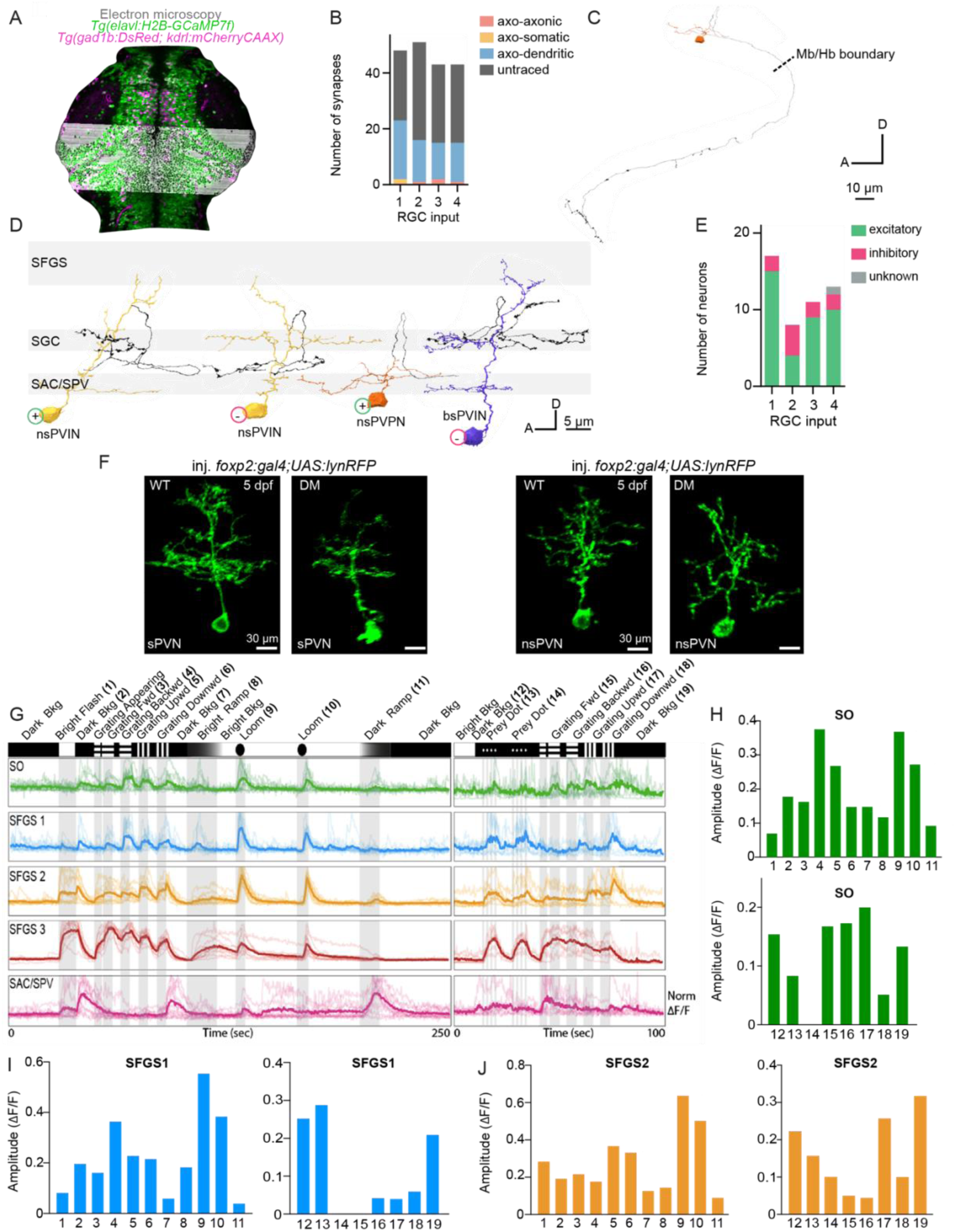
(A) Dorsal view of a 7 dpf *Tg*(*elavl:H2B-GCaMP7f, flk1-mCherryCAAX, gad1B:DsRed*) larval brain showing alignment between two-photon imaging and serial electron microscopy volume. Only the bottom half of the optic tectum is included in this dataset, which spans from the middle of the OT to Rhombomere 3. (**B)** Bar plot showing the distribution of synapse types. **(C)** Detailed morphology of a SAC/SPV-RGC post-synaptic neuron reconstructed from an axo-dendritic synapse. **(D)** Detailed morphology of the most represented SAC/SPV-RGC post-synaptic neurons: nsPVIN, nsPVPN, bsPVIN. **+** and **–** symbols indicate their excitatory or inhibitory nature. Dendrites are color-coded depending on neuronal class, while axons are depicted in black. The nsPVPN neuron labelled in orange is also depicted in (C) in a different orientation. The most superficial OT layers are not shown as none of the post-synaptic neurons innervated them. **(E)** Bar plot showing the distribution of excitatory, inhibitory, and unknown nature of the SAC/SPV-RGC post-synaptic neurons identified. **(F)** Confocal imaging of 5 dpf exemplary stratified (sPVNs) and non-stratified (nsPVNs) in WT and DM larvae, injected at 1-cell stage with a *foxp2:gal4* plasmid. **(G)** Functional responses from the distinct tectal regions to a series of visual stimuli (illustrated above the traces). Data are averaged across larvae; each larva’s trace is represented in shaded color (n = 11 for the first stimulus set; n = 7 for the second). **(H)** Quantification of normalized response amplitudes (ΔF/F) for each stimulus in the SO region, corresponding to the stimuli shown in (G). **(I)** Quantification of normalized response amplitudes (ΔF/F) for each stimulus in the SFGS 1 region, corresponding to the stimuli shown in (G). **(J)** Quantification of normalized response amplitudes (ΔF/F) for each stimulus in the SFGS 2 region, corresponding to the stimuli shown in (G). Scalebars are specified in each panel. Abbreviations: EM, electron microscopy; RGC, retina ganglion cell, SFGS, stratum fibrosum et griseum superficiale; SGC, stratum griseum centrale; SO, stratum opticum; SPV, stratum periventriculare; PVIN, periventricular interneuron; PVPN, periventricular projection neuron, nsPVIN: non-stratified PVIN; nsPVPN, non-stratified PVPN; ms, mono stratified; bs, bistratified; ts, tristratified; s, stratified.

**Supplementary Fig. 6.**
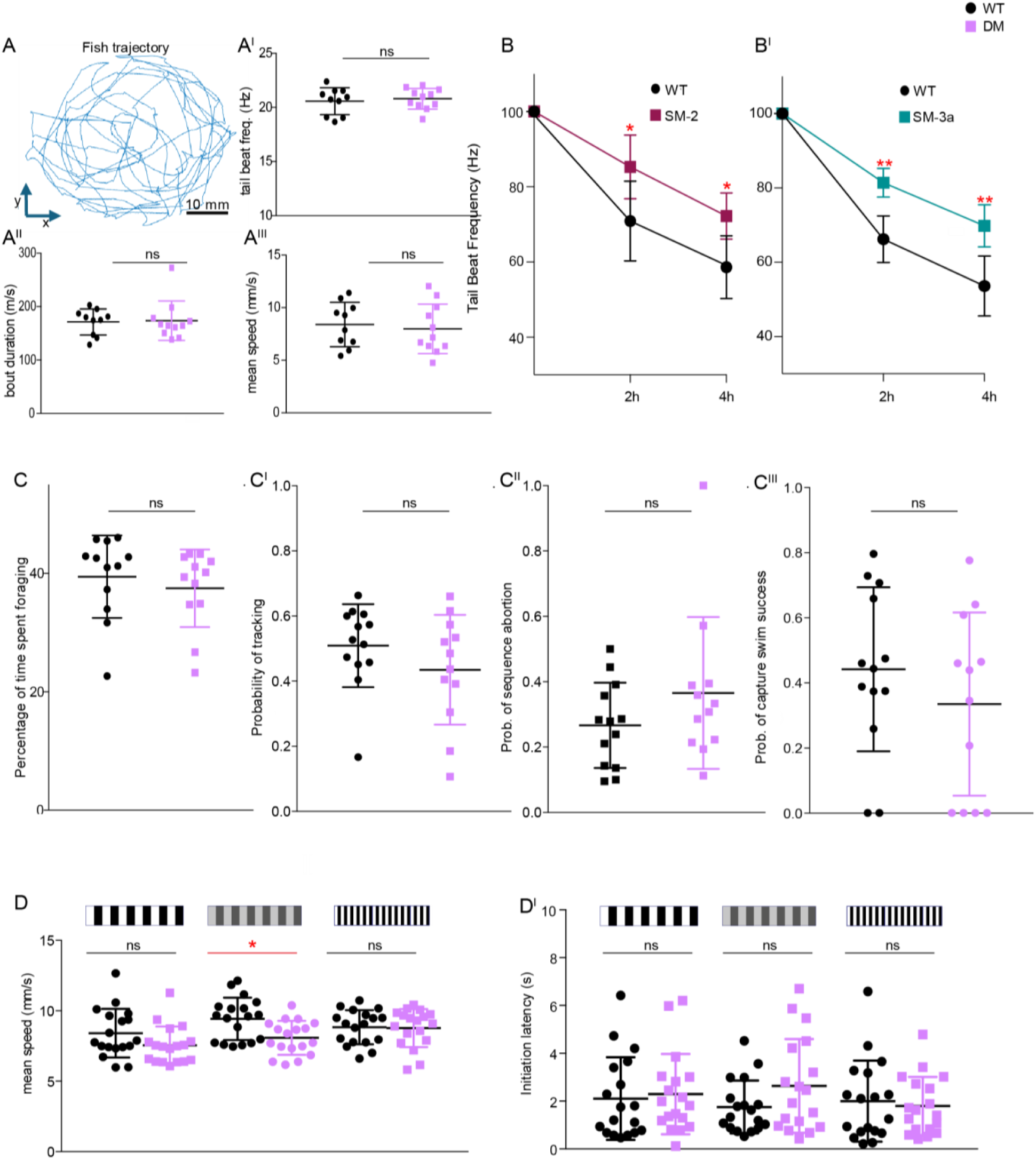
(A-A^III^) Tracking of 6 dpf larvae freely swimming in a petri dish does not show any difference between WT and DM. Representative parameters are presented for the DM larvae: Tail-Beat Frequency (A^I^), bout duration (A^II^), and mean speed (A^III^). Similar results were observed in all the mutants generated (n = 12 larvae per genotype). **(B)** Prey-consumption assay performed at 6 dpf on WT and SM-2 (B), SM-3a (B^I^), larvae, showing percentage (%) of rotifers counted in the dishes after 2 and 4 h. All the mutant larvae eat significantly less than WT controls (n = 10 larvae per genotype). **(C-C^III^)** Quantification and probabilities of different events of the hunting sequence: (C) the percentage of time spent foraging, (C^I^) probability of tracking, (C^II^) probability of sequence abortion, (C^III^) probability of capture swim success (n = 12 larvae per genotype). **(D-D^I^)** Quantification of different variables during OMR response using high and low contrast intensity, and reduced bar size of 6 dpf WT and DM larvae: (D) mean speed (measured in mm/sec), (D^I^) initiation latency (measured in sec) (n = 18 fish per genotype and condition). Horizontal bars = mean ratios; Vertical Bars = standard deviation. Statistical significance was determined in all cases using a non-parametric Mann-Whitney U-test and depicted as: ns – *p* > 0.05; * *p* < 0.05; ** *p* < 0.01. Abbreviations: WT, wild-type; DM, double mutant *lrrn2^-/-^*; *lrrn3a^-/^*^-^; SM-2, single mutant *lrrn2^-/-^*; SM-3a, single mutant *lrrn3a^-/-^*.

**Supplementary Fig. 7.**
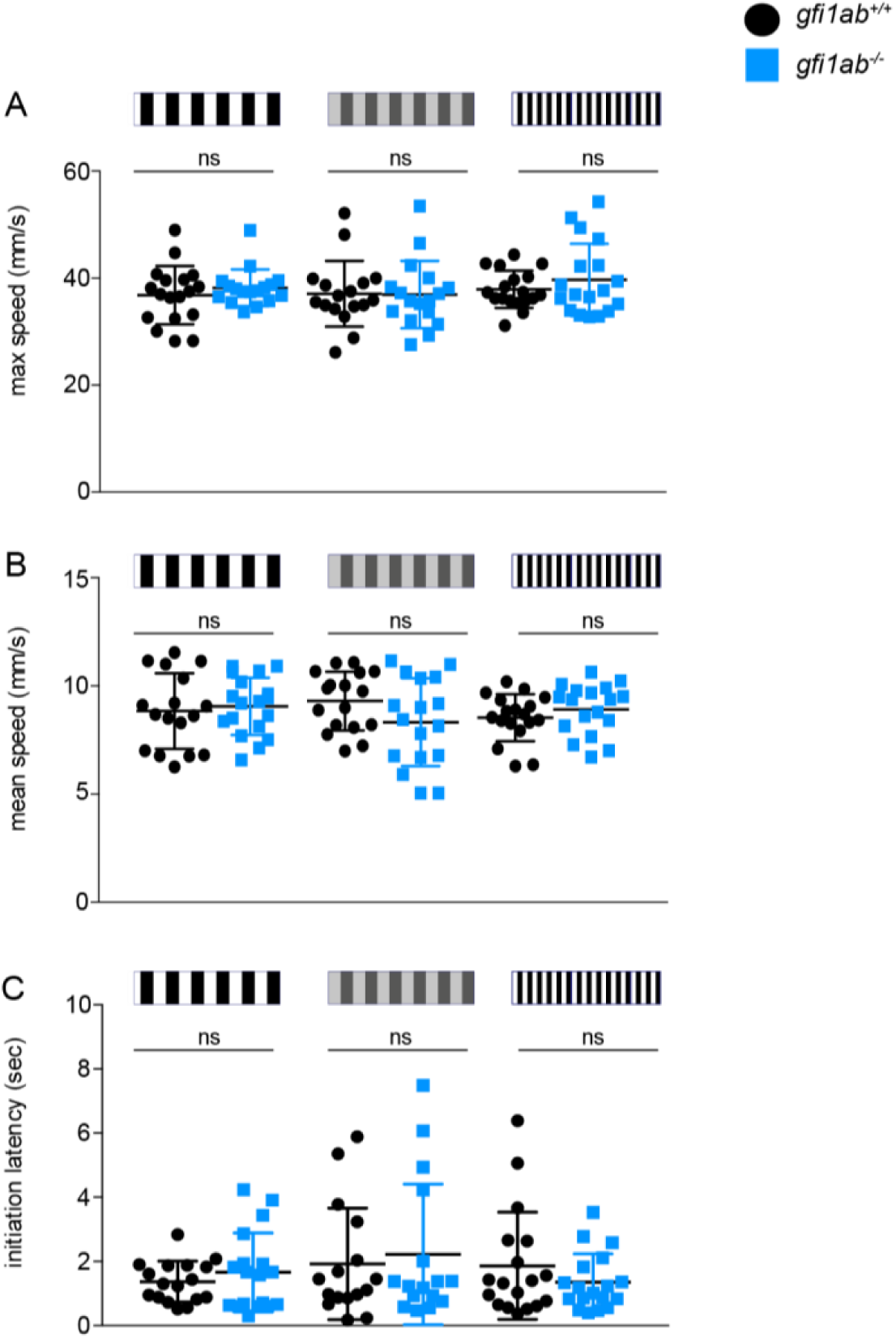
(A) Max speed of 6 dpf WT and *gfi1ab^-/-^*larvae during OMR response using high and low contrast intensity, and reduced bar size measured in mm/sec (WT n = 17 fish, n *gfi1ab^-/-^* = 18 fish per genotype and condition). **(B)** Mean speed of 6 dpf WT and *gfi1ab^-/-^*larvae during OMR response using high and low contrast intensity, and reduced bar size, measured in mm/sec (WT n = 17 fish, n *gfi1ab^-/-^* = 18 fish per genotype and condition). (**C)** Initiation latency of 6 dpf WT and *gfi1ab^-/-^* larvae during OMR response using high and low contrast intensity, and reduced bar size, measured in seconds (WT n = 17 fish, n *gfi1ab^-/-^*= 18 fish per genotype and condition). Horizontal bars = mean ratios; Vertical Bars = standard deviation. Statistical significance was determined in all cases using a non-parametric Mann-Whitney U-test and depicted as: ns – *p* > 0.05.

**Table S1.**
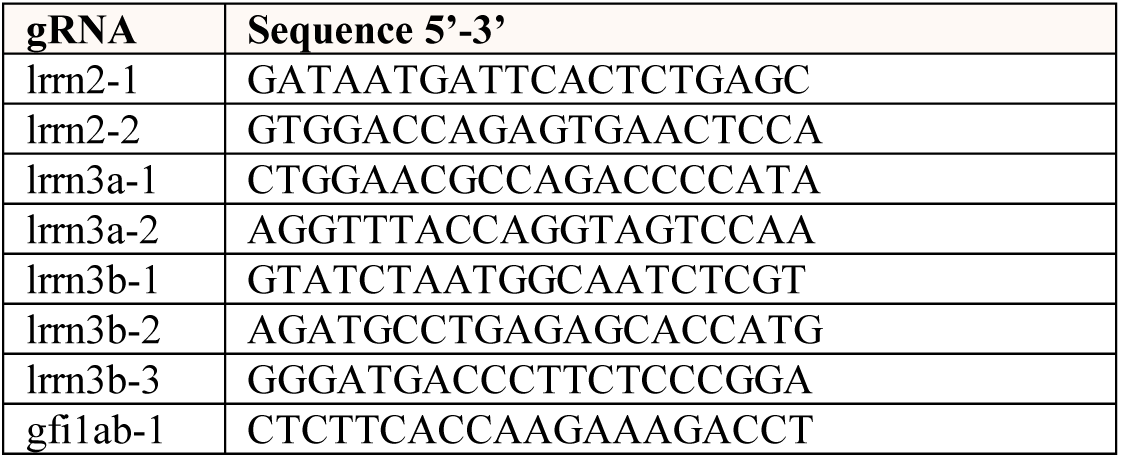

**Table S2.**
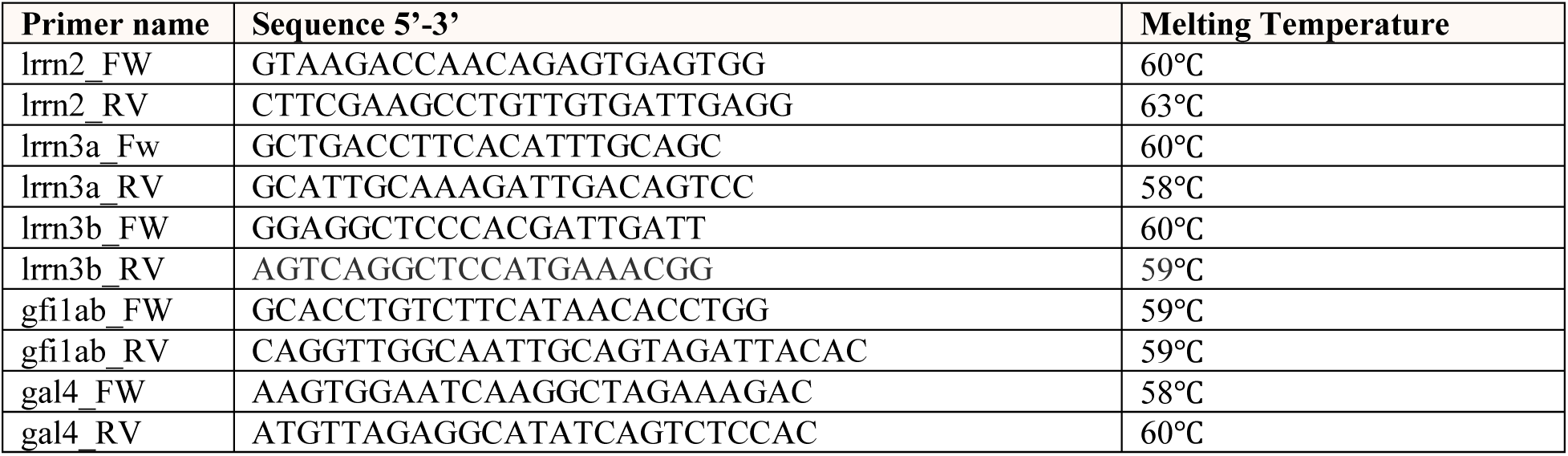

**Table S3.**
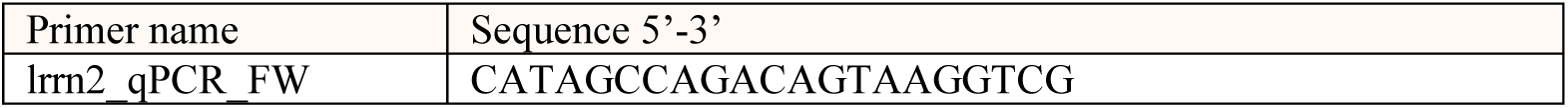
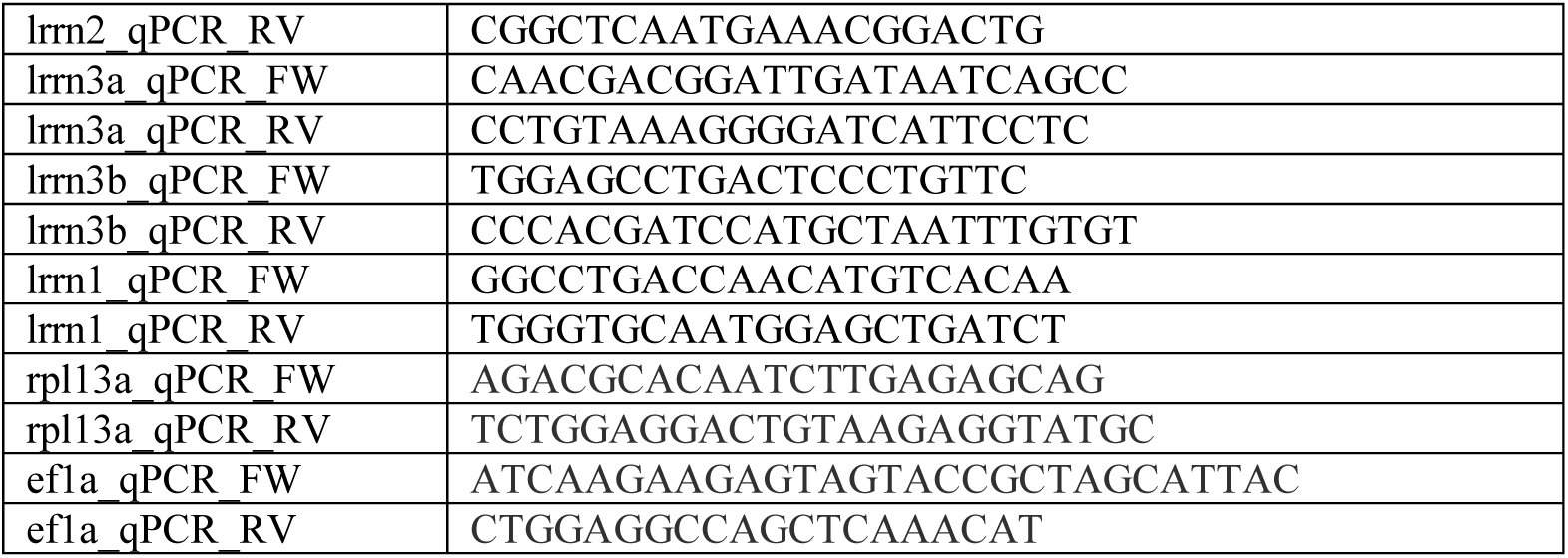

